# Saturation resistance profiling of EGFR variants against tyrosine kinase inhibitors using prime editing

**DOI:** 10.1101/2023.12.03.569825

**Authors:** Younggwang Kim, Hyeong-Cheol Oh, Seungho Lee, Hyongbum Henry Kim

**Author notes:** These authors contributed equally (Younggwang Kim, Hyeong-Cheol Oh, Seungho Lee).

## Abstract

Variants of uncertain significance (VUS) hamper the clinical application of genetic information. For example, in treating lung cancer with tyrosine kinase inhibitors (TKIs), many epidermal growth factor receptor (EGFR) variants remain classified as VUS with respect to TKI sensitivity^1,2^. Such incomplete resistance profiles hinder clinicians from selecting optimal therapeutic agents^3,4^. A high-throughput approach that can evaluate the functional effects of single nucleotide variants (SNVs) could reduce the number of VUS. Here we introduce SynPrime, a method based on prime editing that enabled the generation and functional evaluation of 2,476 SNVs in the *EGFR* gene, including 99% of all possible variants in the canonical tyrosine kinase domain (exons 18 to 21). We determined resistance profiles of 95% (= 1,726/1,817) of all possible EGFR protein variants encoded in the whole tyrosine kinase domain (exons 18 to 24) against afatinib, osimertinib, and osimertinib in the presence of the co-occurring mutation T790M, in PC-9 cells. SynPrime, which uses direct sequencing of endogenous regions to identify SNVs, provided more accurate functional evaluations than a guide RNA abundance-based approach. Our study has the potential to substantially improve the precision of therapeutic choices in clinical settings and contribute to addressing the issue of VUS by being applied to other genes.

## Maintext

Due to recent advances in high-throughput sequencing techniques, a large number of genetic variants are being discovered at an accelerated pace. However, the majority of these genetic variants are classified as variants of uncertain significance (VUS). This lack of information about the functional effects of these variants frequently prevents the determination of optimal clinical treatments in patients with the relevant diseases and the prediction of treatment outcomes.

Targeted therapies using tyrosine kinase inhibitors (TKIs) such as afatinib and osimertinib have been shown to be effective in treating lung cancer with mutations in the epidermal growth factor receptor (*EGFR*) gene^5,6^. These TKIs bind to the intracellular ATP-binding pocket of EGFR in an ATP-competitive manner, preventing ATP from binding and activating the receptor^7^. These TKIs are effective against certain EGFR variants, such as in-frame deletion of exon 19 and L858R^8,9^. However, the effectiveness of these therapies is often compromised by the emergence of drug-resistant variants^10,11^. These TKI-resistant EGFR variants are frequently induced by the introduction of additional substitution mutations^2^. Although some drug-resistant variants have been reported, the majority of EGFR variants remain classified as VUS in relation to TKI resistance. Such variants, which fall into a gray area of clinical actionability, create a conundrum for treatment planning and may lead to suboptimal therapeutic choices^12^. Thus, a comprehensive resistance profile of all possible EGFR variants for representative TKIs like afatinib and osimertinib would provide significant clinical advantages, including more effective TKI selection for individual patients and improved strategies for overcoming drug resistance.

Although the functional effects of variants could be evaluated by expressing them using cDNA-based transgene libraries^13–16^, this non-physiological approach poses a risk of introducing artifacts from overexpression and does not accurately recapitulate the biology driven by these variants in the genomic context. Thus, the generation of variants at endogenous sites could provide more accurate functional evaluation. To introduce many variants into endogenous genes for functional evaluation, various methods such as homology-directed repair (HDR), base editing^17–19^, and prime editing^20^ have been utilized. Although saturation genome editing (SGE) to generate nearly all possible BRCA1 single nucleotide variants (SNVs) by HDR was performed five years ago^21^, this notable approach has not been extended to the SGE of other gene variants, because the efficiency of HDR is usually limited in mammalian cells, requiring haploid cells for accurate functional evaluations. Base editing has been used for high-throughput functional evaluations of variants, owing to its overall high efficiency^22–26^. However, each type of base editor can induce less than 50% of all possible SNVs because such editors i) can only generate C•G to T•A, A•T to G•C, or C•G to G•C mutations and ii) frequently induce bystander editing, falling short of full SGE. Prime editing, which can precisely create any small-sized genetic modification without introducing double-strand breaks^20^, was used to generate and evaluate variants in scattered, limited regions of NPC1 and BRCA2 in a haploidized HEK293 cell line^27^, a cell type less relevant to the pathophysiological function of NPC1 and BRCA2. Although this study is interesting because prime editing was utilized to generate variants at endogenous sites, only 12% to 34% of all possible SNVs (or 13% to 34% of all possible protein variants) in each analyzed exon were generated and evaluated, leaving the remaining 66% to 88% of the variants (66% to 87% of the protein variants) as VUS. In summary, an efficient approach for SGE to generate nearly all possible SNVs at endogenous sites in disease-relevant non-haploid (e.g., diploid) cells for their functional evaluation has yet to be demonstrated.

In this study, we developed a method named SynPrime, which enabled the generation and identification of 2,476 *EGFR* SNVs (1,726 protein variants). These variants include 99% (= 1,137/1,146) of all possible protein variants that can be encoded by SNVs in the canonical tyrosine kinase domain sequence (exons 18 to 21) and 95% (= 1,726/1,817) of all possible protein variants in the whole tyrosine kinase domain sequence (exons 18 to 24)^28^. We then determined complete resistance profiles of these variants against afatinib, osimertinib, and osimertinib in the presence of the co-occurring mutation T790M, using PC-9 cells that contain an in-frame deletion in *EGFR* exon 19, one of the most common EGFR mutations conferring sensitivity to TKIs^8^. By addressing this unmet need for EGFR-related drug resistance profiles, the present study has the potential to substantially improve the precision of therapeutic choices in clinical settings. Furthermore, SynPrime, using which we evaluated nearly all possible variants in a clinically relevant region of EGFR, can be applied to clarify the role of VUS in other genes.

### Sequencing errors can hinder accurate identification of SNVs induced by prime editing

Previous SGE approaches for the generation and evaluation of all possible SNVs in a region of interest have been conducted using haploid cells or cells partially haploidized in that region owing to limited editing efficiencies^21,27,29,30^. However, because most cells in our body are at least diploid, the evaluation of variants in non-haploid cells would provide a more accurate representation of human disease pathophysiology. Furthermore, haploid cells can easily convert to diploid cells and only a very limited range of haploid cell types are available. As a result, the use of haploid cells would substantially restrict generalized applications of SGE. In this study, we employed PC-9, a non-haploid lung adenocarcinoma cell line with a in-frame deletion in *EGFR* exon 19 (E746-A750 deletion), one of the most common *EGFR* mutations conferring sensitivity to TKIs in non-small cell lung cancer (NSCLC)^9^.

The efficiency of SGE is crucial for the efficient and accurate functional analysis of all possible variants. We attempted to utilize prime editing for efficient SGE of *EGFR* mutations and initially used NRCH-PE2max, which has Cas9-NRCH as the nickase domain^31^. We made this choice because SpCas9-NRCH^32^ offers the broadest PAM compatibility among all SpCas9 variants with different PAM compatibilities^33^. We generated PC-9 cells expressing NRCH-PEmax through the transduction of a lentivirus encoding PEmax (Extended Data Fig. 1a). We also designed 1,674 prime editing guide RNAs (pegRNAs) (= 3 pegRNAs/substitution x 3 substitutions/bp x 186 bp) to induce SGE in the 186 bp-long exon 20 of *EGFR* using a preliminary version of DeepPrime-FT, deep learning models that predict pegRNA efficiencies^31^. Based on these designs, we created a lentiviral pegRNA library, named NRCH-exon20, that contains sequences for 1,845 pegRNAs, including the 1,674 pegRNAs mentioned above and other control pegRNAs (Extended Data Fig.1b, Supplementary Tables 1, 2, Methods). To increase the prime editing efficiency, we also lentivirally delivered dominant negative MLH1 (MLH1dn), which inhibits the DNA mismatch repair (MMR) system^34^.

Ten days after the delivery of the NRCH-exon20 library, we performed deep sequencing on exon 20 of endogenous *EGFR*. Deep sequencing revealed that the percentage of reads with a single substitution in the prime-edited cell population was 10%, only slightly higher than 9.2% in the unedited control cell population (Extended Data Fig. 1c). This result suggests that a large portion of the reads with a single substitution in the prime-edited cell population could be attributed to sequencing or PCR errors. To determine the level of prime editing for each substitution, we compared the read counts of each SNV between the prime edited and unedited cells and calculated the odds ratio and *P*-value from Fisher’s exact test. We used odds ratios in addition to *P*-values, because odds ratios, unlike *P*-values, are barely affected by sequencing depth, provided that the sequencing depth is sufficient. We considered that an intended SNV was significantly generated if it met the following criteria: an odds ratio > 3 and *P* < 0.05, and no mutations at the remaining 185 positions. Only 6% (= 34/558) out of a possible 558 (= 3 substitutions/bp x 186 bp) SNVs were identified as being significantly generated (Extended Data Fig. 1d-f). The median frequency of each of these 34 SNVs among all exon 20 sequencing reads was only 0.043% (range: 0.0028% to 0.12%) (Extended Data Fig. 1g). Although these frequencies were higher overall than the median read frequency of 0.011% (range: 0.00050% to 0.13%) for all SNVs, they were too low to accurately identify SNVs by sequencing given that errors from sequencing are about 0.1% per base pair^35,36^.

### SynPrime enables the accurate identification of prime editing-induced SNVs

To enhance the accuracy of SNV identification, we induced one synonymous mutation in addition to the intended mutation as an indicator of prime editing (Fig. 1a). Furthermore, the introduction of this additional one-base pair edit is expected to enhance the efficiency of the original intended editing by minimizing the activity of the MMR system^34^. We also employed the final published version of DeepPrime-FT to identify highly efficient pegRNAs. Because this completed version of DeepPrime-FT predicted that the efficiencies of PEmax would be substantially higher than those of NRCH-PEmax (Fig. 1b), we opted to use PEmax instead of NRCH-PEmax in the subsequent experiments. We generated a lentiviral library, named Syn-exon20, containing sequences for 1,762 pegRNAs that include 1,674 (=186 bp x 3 substitutions/bp x 3 pegRNAs/substitution) pegRNAs intended to induce synonymous substitutions in addition to the intended edit for SGE of *EGFR* exon 20, along with 88 control pegRNAs (Supplementary Tables 1, 2, Methods).

**Figure 1.**
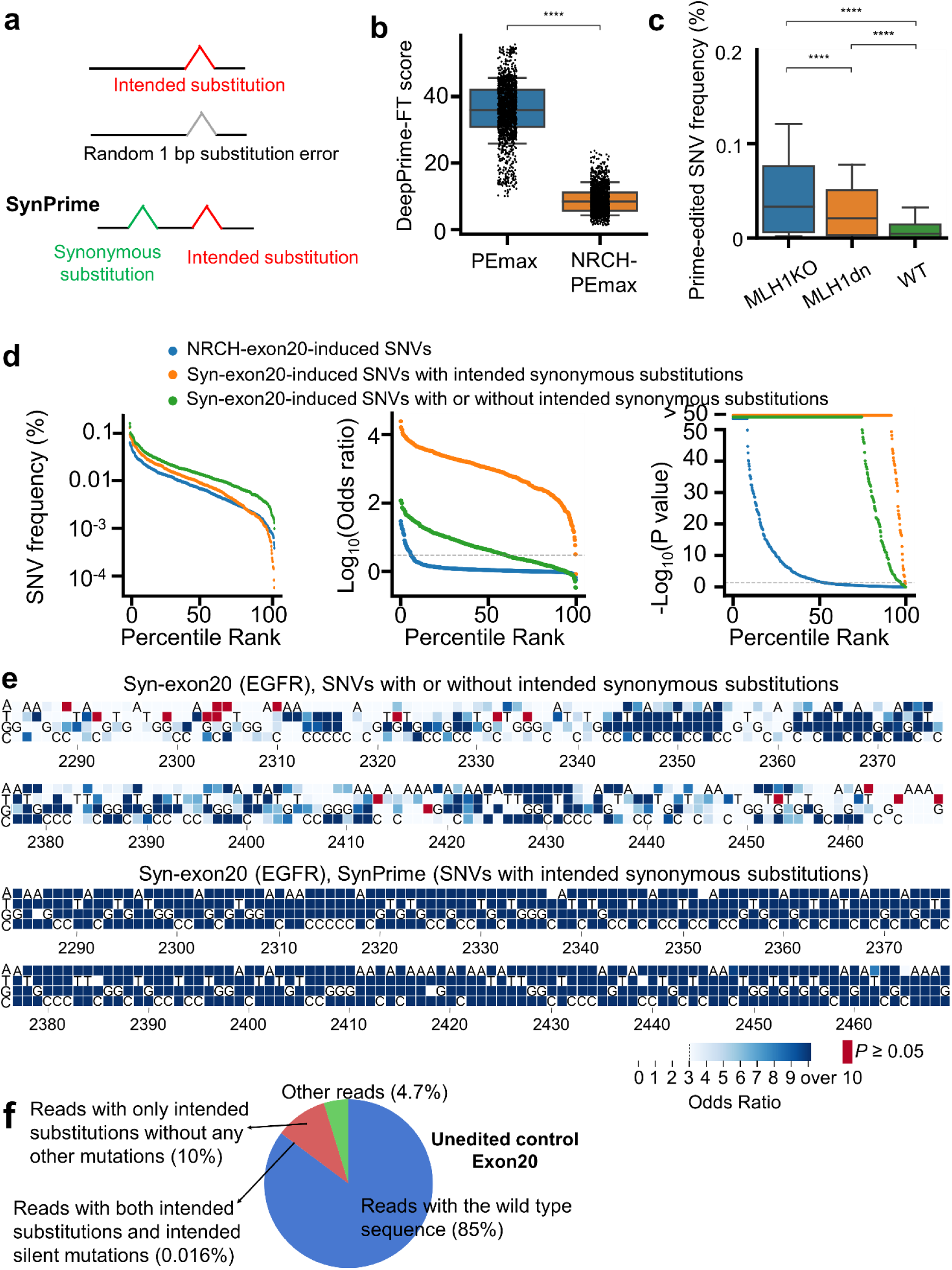
SynPrime enables the accurate identification of prime editing-induced SNVs. **a**, Schematic overview of the SynPrime approach of incorporating an additional synonymous edit near the intended edit. **b**, Distributions of DeepPrime-FT scores for 1,674 pegRNAs for use with PEmax and 1,674 pegRNAs for use with NRCH-PEmax. Boxes represent the 25th, 50^th^, and 75th percentiles, and whiskers show the 10th and 90th percentiles. \*\*\*\**P* <10^-4^ (Two-sided Student’s t-test). **c**, Frequencies of 558 SNVs generated using prime editing in PC-9 cells as a function of the MLH1 status. Boxes represent the 25th, 50^th^, and 75th percentiles, and whiskers show the 10th and 90th percentiles. \*\*\*\**P* <10^-4^ (two-sided paired t-test). **d**, The observed frequencies (left), odds ratios (middle), and *P*-values (right) of 558 SNVs in unedited control cells and prime-edited cells (NRCH-exon20). The dashed lines indicate the positions at which the odds ratio has a value of 3 (middle) and the *P*-value has a value of 0.05 (right). **e,** Heatmap showing odds ratios and/or *P*-values of 558 (= 186 x 3) SNVs generated by prime editing in exon 20 of *EGFR* ten days after the transduction of the Syn-exon20 library. SNVs with *P*-values greater than 0.05 by the two-sided Fisher’s exact test are indicated in red; in these cases, odds ratios are not shown. SNVs with odds ratios lower than 3 are shown in white background. The numbers at the bottom of each heatmap represent the location in the *EGFR* coding sequence. At each position, the nucleotide in the reference sequence is shown. The identification of edited reads is based on the presence of the intended edit regardless of the presence of the intended synonymous substitution (top) or the presence of both the intended edit and the additional synonymous edit (bottom). **f**, Proportion of reads in exon 20 of *EGFR* in unedited control PC-9 cells.

We also postulated that knocking out *MLH1* instead of expressing MLH1dn would lead to higher prime editing efficiencies. We achieved a 99% deletion/indel frequency at the *MLH1* locus in PC-9 cells using SpCas9 and two single-guide RNAs (sgRNAs) targeting *MLH1* (data not shown), suggesting knockout of *MLH1*. Upon evaluating prime editing efficiencies, we found that they were significantly higher in *MLH1*-knockout cells compared both MLH1dn-expressing and wild-type PC-9 cells (Fig. 1c).

Ten days after transducing PEmax-expressing *MLH1*-knockout cells with lentiviral Syn-exon20, we deep-sequenced *EGFR* exon 20. The median frequency of intended SNVs, both with and without intended synonymous substitutions, was 0.038% (range: 0.0014% to 0.36%) (Fig. 1d), which is 3.4-fold (= 0.038%/0.011%) higher than obtained with our initial approach described above. In addition, we also observed a 3.8 (= 4.11/1.08)-fold increase in the median odds ratio and an overall increase in the -log_10_(*P*-values). These increased frequencies of intended SNVs would be attributed to i) the selection of the appropirate PE system (i.e., PEmax instead of NRCH-PEmax), ii) the optimal pegRNA design using the finalized version of DeepPrime-FT, iii) the introduction of additional synonymous mutations, and iv) knocking out *MLH1* as opposed to using MLH1dn. Thanks to this enhanced prime editing efficiency, 60% (= 334/558) out of the 558 (= 3 substitutions/bp x 186 bp) possible SNVs were identified to be significantly generated (Fig. 1d, e top). Although this level is 10-fold (= 60%/6%) higher than the outcome of our initial attempt with NRCH-PEmax, it still falls short of complete SGE, indicating that further improvements are needed for the generation and identification of all possible SNVs.

To improve the accuracy of SNV identification, we utilized an additional synonymous mutation as a marker. When we considered only ’double hits’—sequences with both the intended substitution and the additional synonymous substitution—the median frequency of intended SNVs decreased by 2.5-fold to 0.015% (range: 0.00006% to 0.29%) (Fig. 1d). However, this ’double hit’ counting method, which we have named SynPrime (*Syn*onymous *Prime* Editing), resulted in a 252 (= 1,037/4.11)-fold increase in the median odds ratio and an overall increase in the -log_10_(*P*-values) compared to the approach of counting both ’double hits’ and ’single hits’ (sequences with only the intended substitution without the synonymous substitution). Remarkably, this SynPrime method enabled us to generate and identify 99% (555 out of 558) of all possible SNVs (Fig. 1e), thereby achieving near-complete SGE.

We hypothesized that this substantial improvement in accurate SNV identification using the SynPrime approach could be attributed to effectively filtering out reads from the wild-type allele containing sequencing or PCR errors (i.e., false positive SNV reads). In a control experiment, in which we sequenced the 186-bp long exon 20 of the *EGFR* gene in unedited PC-9 cells, we found that such false-positive SNV reads with one base-pair substitution errors constituted 10% of all reads (Fig. 1f). In stark contrast, when utilizing the SynPrime approach, the frequency of these false-positive reads (i.e., reads with both intended and synonymous substitution errors) in unedited PC-9 cells dropped to just 0.016% of all reads. This value represents a 625-fold reduction (= 10/0.016) compared to results from the method that does not take into account the additional synonymous substitutions.

Additionally, based on the 10% rate of single-point mutations observed in the sequencing of the 186-bp region, we calculated the point mutation error rate in our PCR and sequencing to be approximately 0.06% per base pair. This rate is similar to, or slightly lower than, the generally accepted sequencing error rate of 0.1%^35,36^. We extended our analysis to other regions in unedited control cells and found a tendency for the frequency of false-positive SNVs to increase as the length of the analyzed region grew (Supplementary Fig. 1). Despite this tendency, the calculated frequencies of point mutations per base pair remained consistent across these different regions, ranging from 0.038% to 0.095% with a median value of 0.055%.

### Benchmarking SynPrime-based functional evaluation of SNVs in an essential gene

To evaluate whether our SynPrime approach could be employed for almost saturating functional assessments of SNVs, we prepared a pegRNA library (named Syn-RPL15) that contains 1,492 pegRNA sequences designed to induce 500 SNVs (representing 97% of the 516 total possible SNVs) along with their corresponding synonymous substitutions in the 172-bp long exon 2 of *RPL15*, an essential gene (Supplementary Tables 1,2, Methods). We transduced *MLH1*-knockout PC-9 cells expressing PEmax with the Syn-RPL15 library and conducted puromycin selection to remove untransduced cells, completing this step within five days (Fig. 2a).

**Figure 2.**
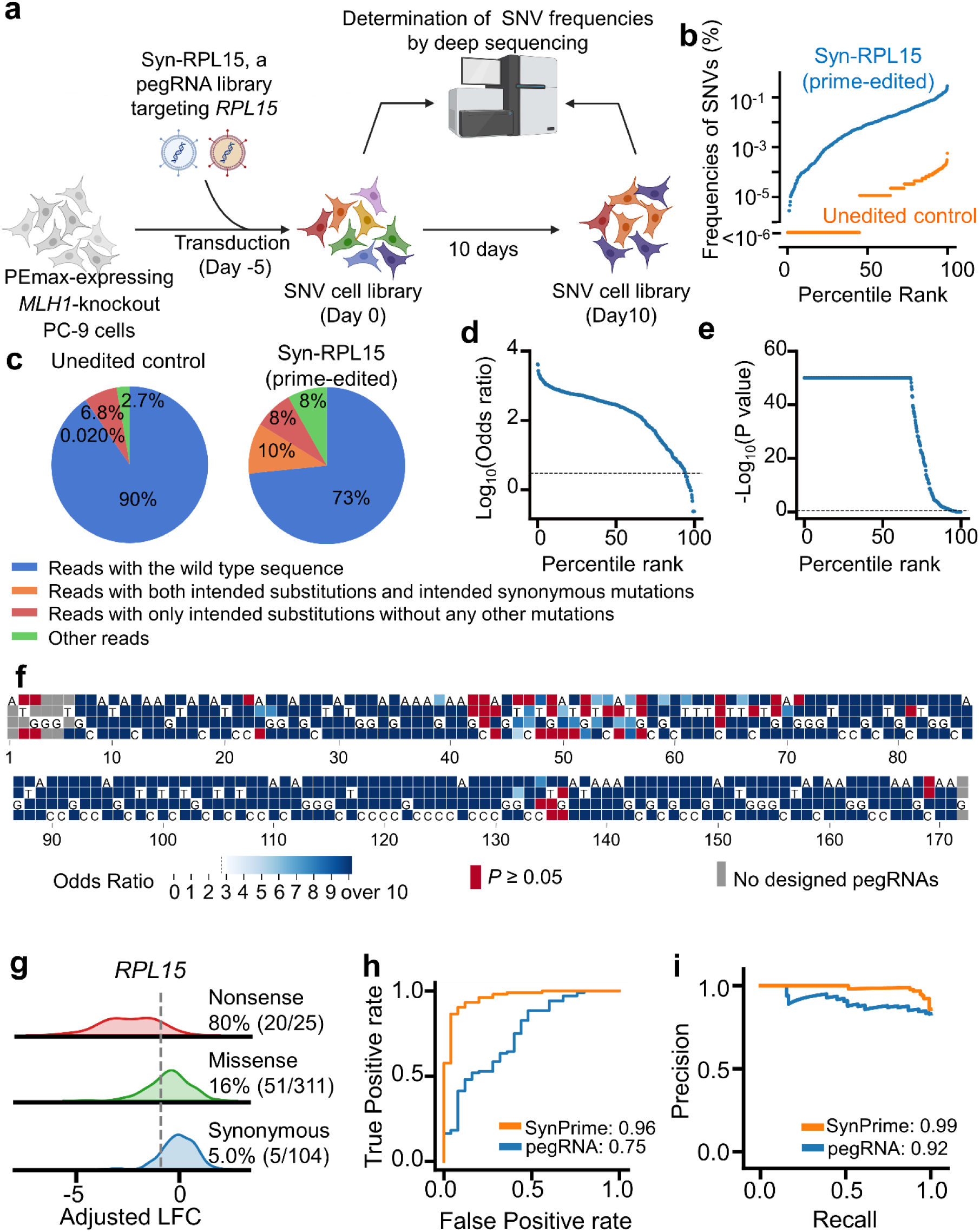
SynPrime evaluation of SNVs in *RPL15*. **a**, Schematic overview of SynPrime high-throughput evaluation of *RPL15* variants. **b**, The observed frequencies of 500 SNVs in unedited control cells and prime-edited cells after treatment with syn-RPL15. **c**, Proportion of sequencing reads containing both intended substitutions and intended synonymous mutations and those containing only intended substitutions without any other mutations in unedited PEmax-expressing PC-9 cells (left, unedited control) and in those cells ten days after transduction with Syn-RPL15 (right). **d**,**e**, The odds ratios (d) and *P*-values (e) of 500 SNVs in PEmax-expressing cells ten days after transduction with Syn-RPL15. The dashed horizontal lines indicate the position at which the odds ratio = 3 (d) and the *P*-value = 0.05 (e). **f**, Heatmap showing odds ratios and/or *P*-values of 516 (= 172 x 3) SNVs generated by prime editing in exon 2 of *RPL15* ten days after the transduction of the Syn-exon20 library. SNVs with *P*-values greater than 0.05 by the two-sided Fisher’s exact test are indicated in red; in these cases, odds ratios are not shown. SNVs with odds ratios lower than 3 are shown in white background. SNVs for which no pegRNAs were designed are shown as gray boxes. The numbers at the bottom of each heatmap represent the location in the *EGFR* coding sequence. At each position, the nucleotide in the reference sequence is shown. **g**, Kernel density estimation plots of adjusted LFCs of SNVs in *RPL15* as a function of the category of SNV. For each category, the number and percentage of SNVs with adjusted LFC values lower than a cutoff value (the gray dashed vertical line), representing the 5th percentile of adjusted LFC values of synonynous mutations, are shown. **h,i,** Receiver operating characteristic (ROC) (**h**) and precision-recall (**i**) curves for adjusted LFCs of SNVs determined by SynPrime (orange) and analysis based on pegRNA abundance (blue) for sets of nonsense (the number of SNVs *n* = 25) versus synonymous SNVs (*n* = 104) in exon 2 of *RPL15*. Area under curve values are shown.

Upon deep sequencing these cells on day 0 (five days post-transduction), we observed that the median frequency of double-hit SNVs (sequences carrying both the intended substitution and the additional synonymous substitution) stood at 0.0083%, with a range from 0.00% to 0.28%. This outcome was a striking 464-fold increase over the median frequency of 1.8 x 10^-5^% observed in unedited cells (Fig. 2b). The sums of these double-hit SNV reads in the prime-edited cell populations and unedited control cells were 10% and 0.02%, respectively, of the total reads (Fig. 2c), indicating that the frequency of identified SNVs was 500-fold (= 10%/0.02%) higher than that of the false-positive SNVs. This double-hit approach significantly identified 440 SNVs, which is 88% (= 440/500) of the designed SNVs and 85% (= 440/516) of all possible SNVs (Fig. 2d, e, f).

These prime-edited cells were cultured for an additional 10 days, and log_2_-fold changes (LFCs) in the frequencies of each SNV at day 10 were calculated relative to those at day 0 (i.e., 5 days after the transduction of Syn-RPL15) to assess the extent of RPL15 function loss associated with the SNV (Fig. 2a). Positional biases in LFCs of synonymous SNVs were modelled using LOWESS (Locally Weighted Scatterplot Smoothing) regression and were used to normalize LFCs by subtraction^21^ (Extended Data Figure 2a, Methods). This normalization slightly increased an area under ROC (AUROC) from 0.92 to 0.96 (Extended Data Fig. 2b). These adjusted LFC (for brevity, LFC) values were well correlated between replicates (Extended Data Fig. 2c). As expected, SNVs causing nonsense mutations were significantly depleted as compared with those causing synonymous mutations (*P* = 1.5 X10^-23^, Student’s t-test, Fig. 2g). We performed Receiver-Operator Characteristics (ROC) and Precision-Recall curve (PRC) analyses, assuming that nonsense SNVs eliminate RPL15 function (thus, the depletion of cells containing these SNVs) and those with synonymous mutations have intact RPL15 function. Our prime editing-based high-throughput evaluations effectively distinguished between loss-of-function and intact RPL15 function, achieving an AUROC of 0.96 and an area under the PRC curve (AUPRC) of 0.99 (Fig. 2h, i), suggesting the high accuracy of our SynPrime approach.

We functionally classified 71 (16% of 440) SNVs, which encode 62 (20% of 307) protein variants, as depleting (i.e., causing loss of RPL15 function, with LFC values < the 5th percentile of the LFC values of synonymous mutations), 189 (43%) SNVs, which encode 178 (58%) protein variants, as non-depleting (i.e., associated with intact RPL15 function, with LFCs > the 20th percentile of the LFCs of synonymous mutations), and the remaining 180 (41%) SNVs, which encode 67 (22%) protein variants, as intermediate (with LFCs between the 5th and 20th percentiles of the LFCs of synonymous mutations) (Extended Data Fig. 2d, e). In summary, using the SynPrime approach, we functionally classified 440 SNVs (85% of all possible SNVs), which encode 307 protein variants, making up 85% of all possible protein variants encoded in exon 2 of *RPL15* (Supplementary Table 3).

### Accuracy of SynPrime vs. pegRNA abundance analyses in *RPL15*

Instead of evaluating endogenous sites, we could calculate the LFCs of lentivirally integrated pegRNA sequences as we and others have previously done for high-throughput functional evaluations^24,25,37–42^. We calculated the LFCs for each SNV using data obtained with the three corresponding pegRNAs, as we have previously done^24^, using the MAGeCK algorithm^43^ (Supplementary Table 3). We observed a high correlation between LFCs of SNV frequencies calculated from pegRNA abundance data from two replicates (Pearson r = 0.81, Extended Data Fig. 2f). As expected, LFC values of pegRNAs for nonsense SNVs were significantly lower than those for synonymous SNVs (*P* = 2.2 X 10^-7^, Student’s t-test, Extended Data Fig. 2g). However, this LFC reduction as assessed by pegRNA abundance-based analysis was less clear than that by the SynPrime approach based on direct sequencing of the endogenous sites (Extended Data Fig. 2g, Fig. 2f); only 32% of nonsense SNVs were slightly depleted by pegRNA abundance-based analysis, whereas 80% were strongly depleted by the SynPrime analysis, raising the possibility that the SynPrime approach would be more accurate and less error-prone than pegRNA abundance-based analysis.

We also conducted ROC and PRC analyses, assuming that pegRNAs that would induce nonsense substitutions would result in the loss of RPL15 function and that negative control pegRNAs, which induce synonymous prime editing, would not affect RPL15 function. We found that AUROC and AUPRC values were 0.75 and 0.92, respectively (Fig. 2h, i), which were lower than the AUROC and AUPRC values obtained from the SynPrime approach described above. We observed only a modest correlation between LFCs determined by endogenous site sequencing (SynPrime) vs. those determined by pegRNA abundance sequencing (r = 0.29, Extended Data Fig. 2h). Taken together, these results suggest that the SynPrime approach is more accurate than pegRNA abundance-based analyses, implying that direct sequencing of endogenous sites can lead to more accurate functional evaluation of the variants.

### Saturating SNV generation in the exons encoding the EGFR tyrosine kinase domain

We next applied SynPrime to determine the effects of saturating point mutations in the EGFR tyrosine kinase domain on drug resistance. This tyrosine kinase domain, which contains 290 amino acids (i.e., requiring 870 coding sequences), is encoded by seven exons (exons 18-24) (Fig. 3a). Using DeepPrime-FT^31^, we designed a total of 7,488 (= 2,515 substitutions x 2∼3 pegRNAs/substitution) pegRNAs that would induce 96% (= 2,515/2610) of all possible SNVs, which correspond to 97% (=1754/1817) of all possible protein variants, in the tyrosine kinase domain. We generated one pegRNA library per exon; these seven libraries were respectively named Syn-exon18 through to Syn-exon24. We also included negative control (i.e., sham-editing or nontargeting) pegRNAs, which comprised about 5% of the pegRNAs in each of the seven libraries (Supplementary Tables 1,2, Methods).

**Figure 3.**
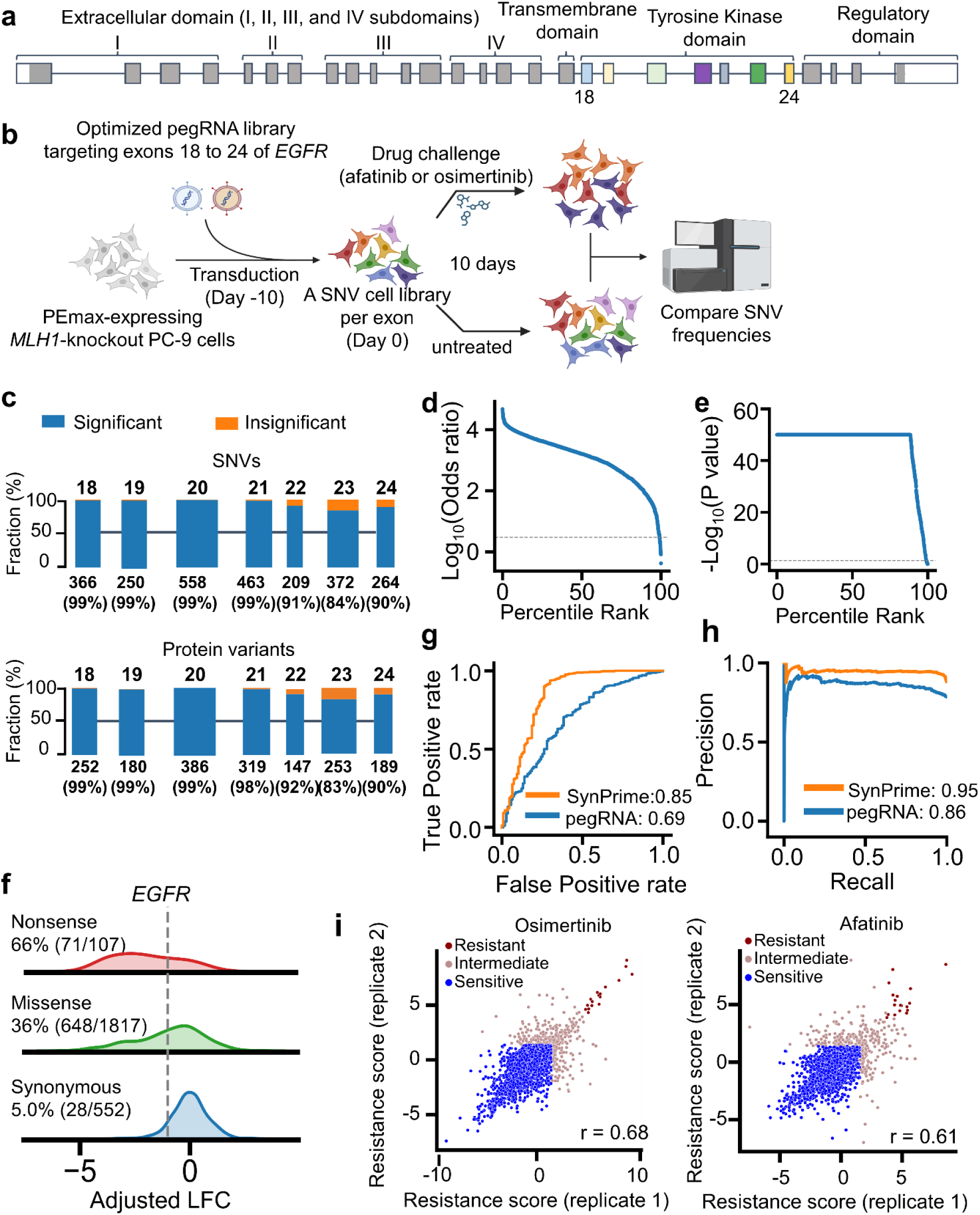
Saturating SNV generation in the region encoding the EGFR tyrosine kinase domain. **a**, Schematic representation of *EGFR*. The tyrosine kinase domain is encoded by exons 18 – 24. The extracellular domain consists of subdomains I, II, III, and IV. **b**, Schematic overview of the functional assay using pooled pegRNA libraries to generate SNVs in *EGFR*. One pegRNA library per exon was created (i.e., a total of seven libraries for seven *EGFR* exons). **c,** The number of significantly generated and identified SNVs (**top**) and protein variants (**bottom**) in each *EGFR* exon. SNVs with frequencies > 0.0005%, odds ratios > 3, and *P*-values < 0.05 at day 0 were considered to be significantly generated. The percentages of significantly generated SNVs (top) and protein variants (bottom) are shown in parentheses (e.g., 99% of the SNVs for exon 18). **d,e**, The odds ratios (d) and *P*-values (e) of 2,515 SNVs in PEmax-expressing cells ten days after transduction with Syn-exon18, Syn-exon19, …, and Syn-exon24. The dashed horizontal lines indicate the position at which the odds ratio = 3 (d) and the *P*-value = 0.05 (e). **f**, Kernel density estimation plots of adjusted LFCs of SNVs in the region encoding the EGFR tyrosine kinase domain as a function of the category of SNV. For each category, the number and percentage of SNVs with adjusted LFC values lower than a cutoff value (the gray dashed vertical line), representing the 5th percentile of adjusted LFC values of synoynous mutations, are shown. **g,h,** Receiver operating characteristic (ROC) (**g**) and precision-recall (**h**) curves for adjusted LFCs of SNVs determined by SynPrime (orange) and analysis based on pegRNA abundance (blue) for sets of nonsense (the number of SNVs *n* = 107) versus synonymous SNVs (*n* = 552) in exons 18-24 of *EGFR*. Area under curve values are shown. **i**, Correlation between SynPrime LFC values following treatment with afatinib (left) and osimertinib (right) in two biological replicates. The classification of each SNV is indicated by the dot color. Pearson correlation coefficients are shown. The number of SNVs n = 2,476 (osimertinib), 2,476 (afatinib).

We induced SGE by independently transducing each library into PEmax-expressing *MLH1* knockout PC-9 cells, after which we cultured them for 10 days to induce sufficient prime editing (Figure 3b). Deep sequencing revealed that the sums of the two-hit SNV reads (i.e., reads with the intended substitution and the additional intended synonymous mutation) in the unedited control and prime-edited cell populations were a median of 0.009% and 19%, respectively, of the total reads (Supplementary Fig. 2), suggesting that the frequency of identified SNVs was a median of 2,111-fold (= 19%/0.0009%) higher than the frequency of false positive PCR and sequencing errors. We filtered out 39 non-significant SNVs (i.e., odds ratio ≤ 3 or *P*-value ≥ 0.05 in any replicate) and identified the remaining 2,476 significant SNVs (95% of all 2,610 possible SNVs), which correspond to 1,726 protein variants (95% of all 1,817 possible protein variants) (Fig. 3c-e, Extended Data Fig. 3, Supplementary Table 4).

### Accuracy of SynPrime vs. pegRNA abundance analyses in *EGFR*

When we evaluate the SNV libraries at day 10 (20 days after the transduction of pegRNA libraries), SNVs causing nonsense mutations were significantly depleted as compared with those causing synonymous mutations (*P* = 1.8 X 10^-67^, Student’s t-test, Fig. 3f), suggesting the dependence of PC-9 cells on EGFR signaling. However, by pegRNA abundance-based analysis, the depletion of nonsense SNVs was less prominent than that by the SynPrime (Supplementary Fig. 3a, b). AUROC and AUPRC values of the SynPrime approach were 0.85 and 0.95, respectively, whereas those of the pegRNA abundance-based analyses were 0.69 and 0.86, respectively (Fig. 3g, h). We also observed only a modest correlation between LFCs determined by endogenous site sequencing (SynPrime) vs. pegRNA abundance sequencing (r = 0.40, Supplementary Fig. 3c) as similarly described above for *RPL15*. Taken together, these results corroborate that the SynPrime approach is more accurate than pegRNA abundance-based analyses.

### Complete resistance profiles of 2,476 *EGFR* SNVs against afatinib and osimertinib

We cultured the SNV libraries for an additional 10 days under three different experimental conditions: the control condition without any treatment, treatment with the 2^nd^ generation TKI afatinib, and treatment with the 3^rd^ generation TKI osimertinib (Figure 3b). LFCs of SNV frequencies in the TKI-treated arms were compared with those in the untreated arm to determine their effect on resistance to these two TKIs (Supplementary Table 4, Methods). We named these normalized LFCs resistance scores. The resistance scores of SNVs in both the osimertinib and afatinib arms from two different replicates correlated well (Fig. 3i). Based on the resistance scores, we classified SNVs into “resistant” (the resistance score exceeded the resistance score of synonymous SNVs in the 99.7^th^ percentile in both replicates), “sensitive” (the resistance score was lower than the resistance score of synonymous SNVs in the 95th percentile in both replicates), and “intermediate” (the remaining SNVs) categories.

Out of the 1,726 protein variants, 158 were encoded by two SNVs and 20 were encoded by three SNVs. When we evaluated the correlations between resistance scores of the two members of each pair (note that each trio comprises three pairs), we observed high correlations between them (Supplementary Fig. 4a, left and middle). We then used the mean of SNV resistance scores to calculate the resistance score of each protein variant. These resistance scores of protein variants in both the osimertinib and afatinib arms from two different replicates correlated well (Supplementary Fig. 4b, left and middle). Regarding afatinib, 2,138, 320, and 18 SNVs, which encode 1476, 260, and 16 protein variants, were respectively classified as sensitive, intermediate, and resistant (Extended Data Fig. 4, Supplementary Fig. 5). Fourteen (88%) of the 16 resistant protein variants have not been previously reported (Supplementary Table 5, Supplementary Fig. 4c). Regarding osimertinib, 2,102, 357, and 17 SNVs, which encode 1,448, 290, and 15 protein variants, were respectively classified as sensitive, intermediate, and resistant (Extended Data Fig. 5, Supplementary Fig. 6). Twelve (80%) of the 15 resistant protein variants have not been previously reported (Supplementary Table 5, Supplementary Fig. 4c).

### Resistance profiles of 2,391 *EGFR* SNVs against osimertinib in PC-9-T790M cells

T790M is the most common gatekeeper mutation that induces resistance to 1^st^ and 2^nd^ generation TKIs^10,44^. Given that osimertinib is the first-line treatment choice in patients with T790M-positive advanced NSCLC^45^, we next determined the resistance profiles of SNVs against osimertinib in PC-9 cells containing T790M. We introduced the T790M (c.2369C>T) mutation into the PEmax-expressing PC-9 cells line by using cytosine base editing, generating PC-9-T790M (Supplementary Fig. 7a-c, Methods). Sanger sequencing and deep sequencing showed that the frequency of alleles containing T790M was 99.8% (Supplementary Fig. 7d, data not shown), suggesting that the T790M mutation was homozygous. We conducted SynPrime evaluation in these PC-9-T790M cells using osimertinib given that the T790M mutation confers resistance to afatinib (Supplementary Table 4). Resistance scores of SNVs and protein variants in the PC-9-T790M cells were highly correlated between two replicates (Supplementary Fig. 7e, 4b). We also observed high correlations between resistance scores of the members of each pair of SNVs encoding identical protein variants (Supplementary Fig. 4a, right), an observation that held true for both the afatinib and osimertinib arms. The resistance scores of protein variants from two different replicates also correlated well (Supplementary Fig. 4b, right). We classified 2,028, 335, and 28 SNVs, which encode 1,395, 281, and 26 protein variants, as sensitive, intermediate, and resistant, respectively (Extended Data Fig. 6, Supplementary Fig. 7). Eighteen (69%) of the 26 protein variants causing resistance have not been previously reported (Supplementary Table 5, Supplementary Fig. 4c).

### Resistance profiles of *EGFR* SNVs against afatinib and osimertinib in the absence of T790M and against osimertinib in the presence of T790M

We compared the resistance profiles of SNVs in the region encoding the EGFR tyrosine kinase domain for three conditions (afatinib and osimertinib in the absence of T790M and osimertinib in the presence of T790M) in PC-9 cells, providing an unprecedented list of resistance profiles against afatinib and osimertinib for almost all possible SNVs (Figure 4, Extended Data Fig. 4-6, Supplementary Fig. 5,6, 8, Supplementary Table 4). Notable findings are as follows.

**Figure 4.**
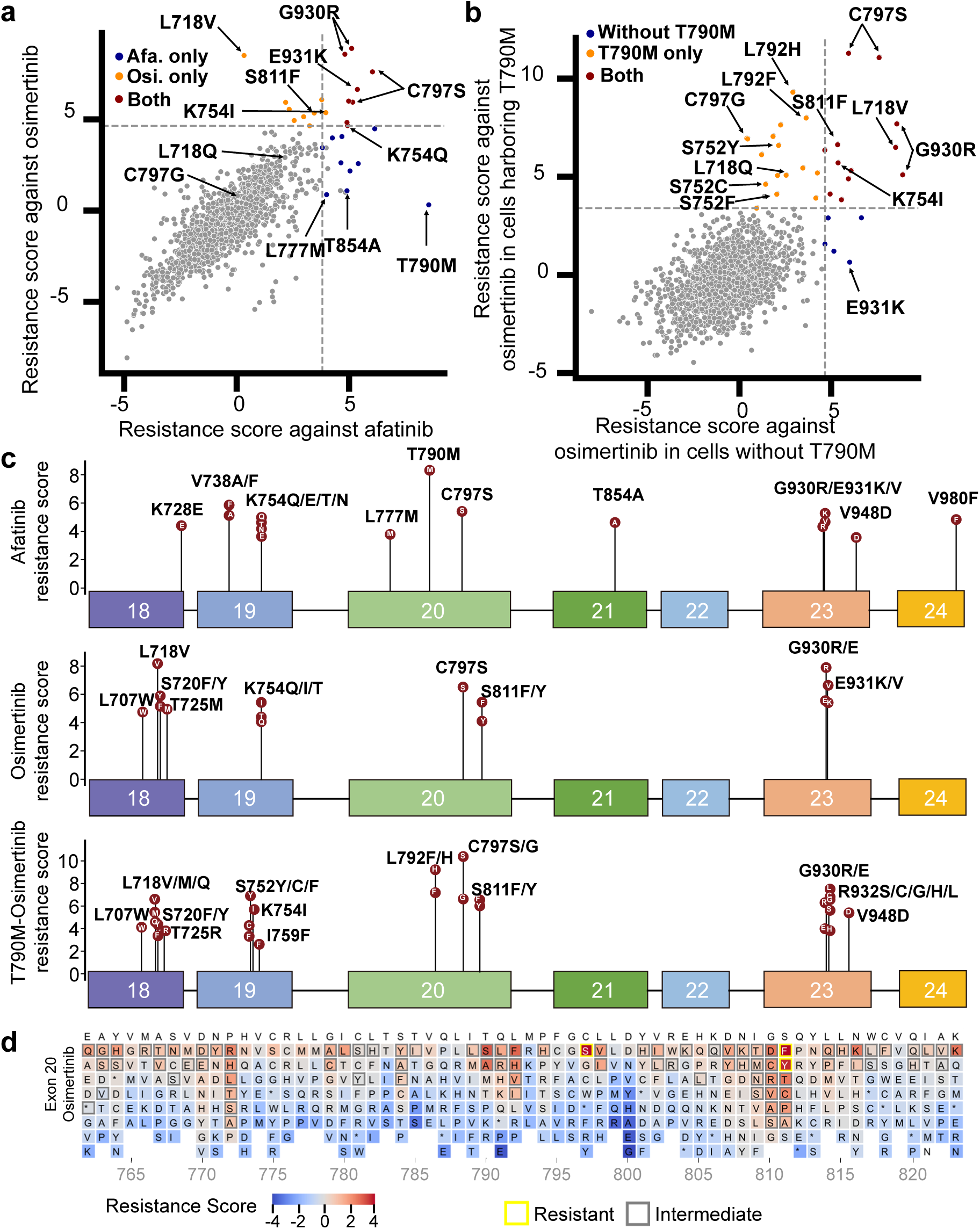
Landscape of TKI resistance conferred by SNVs in the region encoding the EGFR tyrosine kinase domain. **a**, Resistance scores of *EGFR* SNVs in PC-9 cells lacking the T790M gatekeeper mutation (a) and against osimertinib in the presence (the y axis) and absence (the x axis) of the T790M mutation in PC-9 cells (b). The dashed lines represent the lowest resistance score values among the resistant SNVs for each evaluation. SNVs that confer resistance against any of TKIs are shown in orange, red, or blue depending on the TKI or the presence of co-occuring T790M mutation. Afa., afatinib; Osi., osimertinib. **c**. Lollipop plot of SNVs in *EGFR* exons 18 to 24 and their resistance scores against afatinib (top), osimertinib (middle), and osimertinib in the presence of a co-occuring T790M mutation (bottom). For brevity, only SNVs classified as conferring resistance are plotted. **d**, Heatmap showing osimertinib resistance scores of 1,817 protein variants. These variants were generated by prime editing in exon 20 of *EGFR* in PC-9 cells. Boxes outlined in yellow and gray indicate protein variants that confer resistant and intermediate phenotypes, respectively. The numbers at the bottom of each heatmap represent the location in the EGFR amino acid sequence. At each position, the amino acid in the reference sequence is shown at the top. Protein variants with *P*-values greater than 0.05 or odds ratios lower than 3 were excluded from the analysis and are shown in white background.

T790M was classified as resistant in the afatinib arm but not in the osimertinib arm (Fig. 4a), which is in line with a previous report that T790M is the most common gatekeeper mutation that induces resistance to 1^st^ and 2^nd^ generation TKIs^10,44^. C797S, a previously well-characterized mutation conferring resistance to both osimertinib and afatinib^11,46,47^, was classified as resistant in all three arms. Taken together, the results of our SynPrime evaluation are compatible with representative previous reports about TKI resistance profiles.

Interestingly, we found that nine protein variants including L718V caused resistance to osimertinib but not afatinib, suggesting that we could use afatinib to treat patients with this mutation. Compatible with our results, we found a case report suggesting that the L718V mutation confers resistance to osimertinib, but not afatinib^48^. We also found that L718Q caused resistance to osimertinib in PC-9-T790M cells and that the effect was intermediate to that of afatinib and osimertinib in PC-9 cells. This result is in line with a previous report that L718Q causes osimertinib resistance in NSCLC regardless of the presence of T790M, possibly due to spatial restrictions and reduced hydrophobic interactions of EGFR with osimertinib^49^. Our finding is also compatible with another previous report that afatinib can be a possible treatment option with modest efficacy in patients harboring the L718Q mutation^15,50^.

We also found that 16 protein variants including S752Y/C/F and C797G caused osimertinib resistance only in the presence of T790M. In the absence of T790M, however, these variants were associated with an intermediate or sensitive phenotype for osimertinib, suggesting that osimertinib could be used in patients bearing any one of those 16 protein variants in the absence of T790M. This finding is compatible with a previous report that a SNV encoding S752C was identified after osimertinib treatment in patients with the T790M mutation^51^ and with several reports that have indicated that C797G is associated with osimertinib resistance in the presence of T790M^52,53^. In the absence of T790M, cells carrying C797G were sensitive to both afatinib and osimertinib. Although there have been no reports about the resistance profile of C797G against afatinib and osimertinib in the absence of T790M, our results suggest that both afatinib and osimertinib could be considered for the treatment of patients bearing C797G in the absence of T790M.

L777M and T854A were classified as resistant to afatinib, compatible with previous reports that the two mutations are acquired resistance mutations in response to a 1^st^ generation TKI^54,55^, and as sensitive to osimertinib treatment^56^. Although there are no reports on the resistance profiles of the L777M and T854A mutations in relation to afatinib, a 2^nd^ generation TKI, our SynPrime evaluation suggests that osimertinib could be effective in patients with L777M- or T854A-harboring tumors.

We identified clustered, resistance-conferring SNVs that affected K754 (encoded in exon 19), S811 (exon 20), and G930R and E931 (exon 23). K754I conferred resistance against osimertinib in both the presence and absence of T790M; it was classified as having an intermediate effect in the afatinib arm. This result is compatible with a case report about a patient bearing T790M, in which osimertinib treatment led to loss of the T790M-encoding SNV but enriched that encoding K754I^57^.

Of note, because exons 18-21 encode the region comprising the ATP binding pocket, current clinical tests for EGFR mutations mainly focus only on these four exons^58^. The region encoded by exons 22-24, outside of the ATP binding pocket^59^ (Fig. 5a), has not been extensively investigated. G930R, encoded in exon 23, was classified as resistant in all three arms; E931K was classified as resistant against both afatinib and osimertinib, but only in the absence of T790M. Previous crystallographic studies revealed that E931 and G930 are located on the periphery of the asymmetrical kinase-activating dimer interface, and may affect the dimerization and activation potential of EGFR^59,60^. Taken together, our results are compatible with previous reports and provide new information about sensitivity to and resistance against TKIs, suggesting possible drug choices for a given EGFR mutation profile.

**Figure 5.**
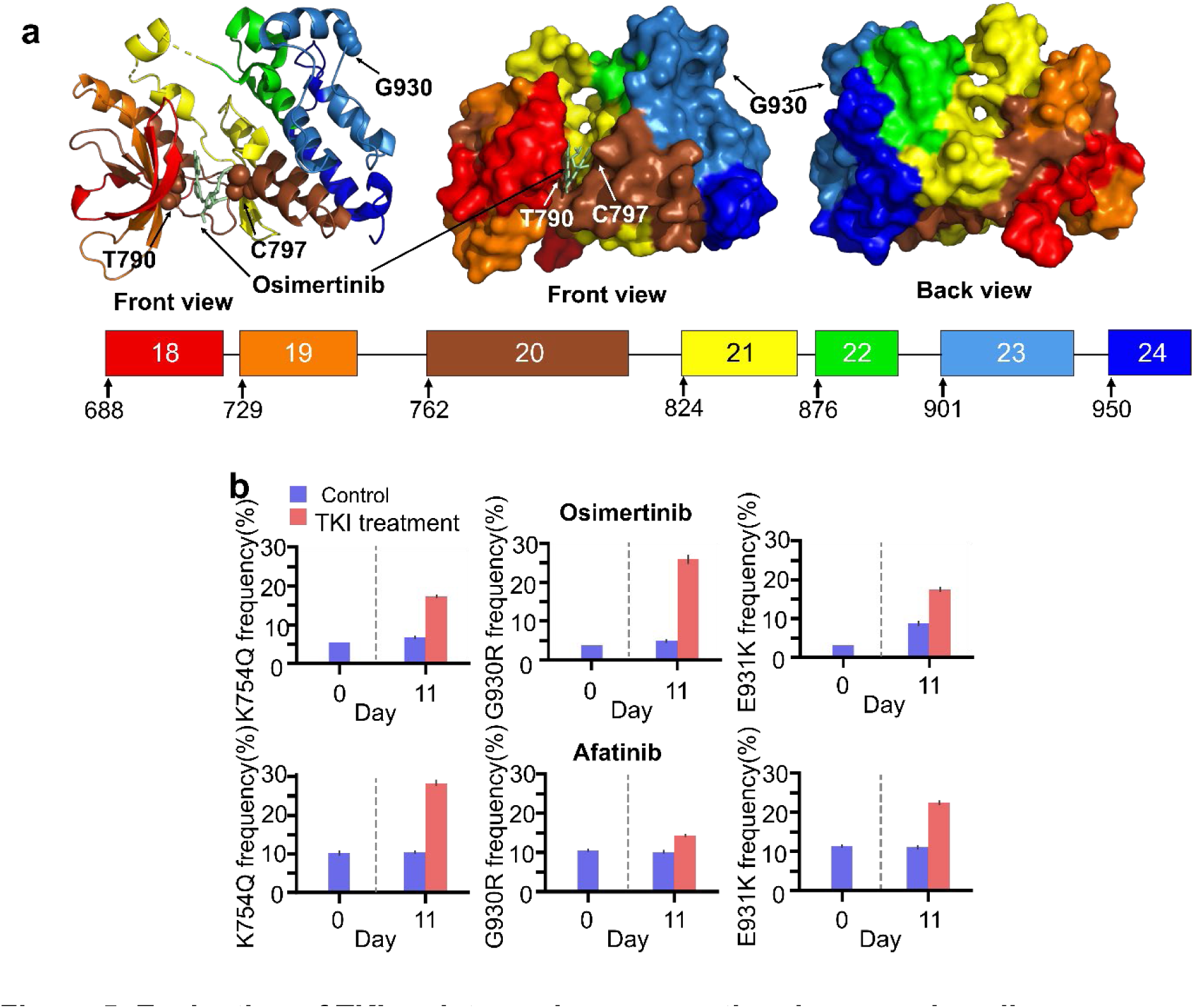
Evaluation of TKI resistance in a conventional manner in cells. **a**, Location of the G930 residue in the crystal structure of the EGFR tyrosine kinase domain bound to osimertinib (Protein data bank: 6JXT, EGFR complex with osimertinib). The amino acid regions corresponding to the coding sequence of each exon are indicated in different colors. **b**, Percentage of reads containing intended edits before (D0, blue) and after (Day 11) treatment with osimertinib (red, day 11) or control solvent (blue, day 11). The number of biological replicates for treatment with osimiertinib (top) and afatinib (bottom) *n* = 3.

### Evaluation of TKI resistance in a conventional cell-based manner

To evaluate the resistance of cells bearing the novel SNVs identified in the current study in a conventional manner, we individually delivered lentivirus encoding pegRNAs that would generate SVNs in *EGFR* encoding K754Q, G930R, or E931K to PEmax-expressing *MLH1*-knockout PC-9 cells (Supplementary Fig. 8). We mixed these prime edited cells with control cells at a 25:75 ratio. These mixed cell populations were then divided into untreated and afatinib- or osimertinib-treated conditions, cultured for 10 days, and subjected to deep-sequencing. Mean frequencies of SNVs encoding K754Q, G930R, and E931K increased 2.6-fold (= 17%/6.7%), 5.4-fold (= 26%/4.8%), and 2.0-fold (= 17%/8.7%), respectively, in the presence of osimertinib and 2.7-fold (= 28%/10%), 1.4-fold (= 14%/10%), and 2-fold (= 22%/11%) in the presence of afatinib, respectively, as compared to the untreated control (Fig. 5a, Supplementary Table 6), corroborating that these SNVs confer resistance to osimertinib and afatinib.

## Discussion

In this study, we evaluated 2,476 (95%) of the SNVs (1726 (95%) of 1817 possible protein variants) that can be generated by single point mutations in the EGFR tyrosine kinase domain. To be used for decision making in clinical settings, drug resistance profiles should be highly accurate. To maximize the accuracy, we i) directly analyzed endogenous sequences instead of relying on analyses of guide RNA abundance^22,23,40,42^ or surrogate reporter targets^24,25,41^, ii) substantially reduced sequencing and PCR-related noise as compared to the signal (i.e., frequencies of intended SNVs) using additional synonymous editing (SynPrime (double-hit approach)), iii) increased prime editing efficiencies using DeepPrime-based pegRNA design, knocking out *MLH1*, and introducing synonymous mutations, to maximize the signal, iv) classified only significantly generated SNVs by filtering out those with either insignificant *P*-values or odds ratios rather than simply filtering out reads with frequencies lower than a certain value^27^, and v) classified SNVs as either sensitive or resistant only when the results from two biological replicates matched.

Because the functional effects of SNVs are the result of mutations introduced into endogenous gene regions, we can easily assume that functional classifications based on direct analyses of endogenous sequences would be more accurate than those based on indirect evaluation of endogenous regions by analysis of guide RNA abundance or surrogate reporter targets. However, direct evidence for this assumption has yet to be provided. In this study, we provide results that indicate that direct analysis of endogenous sites can be more accurate for the functional classification of SNVs using two different genes (i.e., *RPL15* and *EGFR*).

A problem in the approach based on direct analyses of endogenous sites is the very low frequencies of intended SNVs. Given that the measured frequencies of SNVs generated by high-throughput prime editing in an endogenous region are diluted by the size of the pegRNA library used in the study, the identification of intended SNVs from deep sequencing reads can be challenging, especially when the library is relatively large. For example, if we assume that a library contains 600 pegRNA sequences for inducing 600 different intended edits at 200-bp long endogenous sites and that the prime editing efficiency is as high as 60% for a given intended edit, the measured frequency of the intended SNV would be only 0.1% (= 60%/600), which is similar to generally regarded values for deep sequencing errors. If we use a bigger library or if the prime editing efficiency is lower than 60%, which is usual, then the measured SNV frequency would be lower than 0.1%, making it difficult to accurately identify intended SNVs from sequencing errors. Indeed, in our study, the measured median SNV frequencies ranged from 0.009% to 0.012% in the seven *EGFR* exons. To increase the accuracy of intended SNV identification, we added another substitution, which reduced false positive reads by median 626 (= (10% + 0.016%)/0.016%)-fold, increasing the signal to noise ratio by median 443 (= (15%/0.016%)/((15% + 6.2%)/(0.016% + 10%)))-fold.

EGFR mutations other than well-studies mutations (exon 19 in-frame deletion, L858R, T790M, and C797S) account for up to 18% of all EGFR mutations observed in patients^61^. However, a lack of resistance profiles of these diverse mutations against TKIs has hindered the selection of optimal therapeutic agents for individual patients with those mutations. By providing a comprehensive resistance profile for EGFR variants against TKIs, we are stepping closer to truly personalized medicine. In addition, with the applications of SynPrime to other genes, the issues surrounding VUSs can be addressed, possibly ushering in a new era of precision medicine.

## Acknowledgements

We would like to thank Younghye Kim and Gahyun Baek for assisting with the experiments. This work was supported, in part, by the National Research Foundation of Korea grant funded by the Korean government (MSIT) (2022R1A3B1078084 (H.H.K.), 2018R1A5A2025079 (H.H.K.). The Bio and Medical Technology Development Program of the NRF funded by the Korean government (MSIT) (2022M3A9E4017127 (H.H.K.), 2022M3A9F3017506 (H.H.K.) and (RS-2023-00260968) (H.H.K.)), the Korea Drug Development Fund funded by the Ministry of Science and ICT, the Ministry of Trade, Industry and Energy, and the Ministry of Health and Welfare, Republic of Korea (HN21C0917 (H.H.K.)), the Yonsei Signature Research Cluster Program of 2023-22-0012 (H.H.K.), the Brain Korea 21 FOUR Project for Medical Science (Yonsei University College of Medicine), the SNUH Kun-hee Lee Child Cancer and Rare Disease Project, Republic of Korea (22B-000-0101 (H.H.K.)), the Yonsei Fellow Program, funded by Lee Youn Jae, and the Korea Health Technology R&D Project funded by the Ministry of Health and Welfare, Republic of Korea (HI21C1314 (H.H.K.)); Genome editing research program funded from the Korea government (MSIT)(RS-2023-00263285 (Y.K.)), Basic Science Research Program through the National Research Foundation of Korea funded by the Ministry of Education (2022R1I1A1A01066096 (Y.K.)), a faculty research grant of Yonsei University College of Medicine (6-2022-0078(Y.K.)), the financial support of the Catholic Medical Center Research Foundation made in the program year of 2023 (Y.K.); a grant of the MD-PhD/Medical Scientist Training Program (H.C.O.) through the Korea Health Industry Development Institute (KHIDI), funded by the Ministry of Health & Welfare, Republic of Korea.

## Author contributions

Y.K., H.C.O., S.L. and H.H.K. conceived and designed the research. Y.K., H.C.O. and S.L. performed the experiments and data analysis. Y.K., H.C.O., S.L. and H.H.K. wrote the manuscript.

## Online Methods

### Design of the pegRNA libraries

Using the reference human genome (hg38) as a guide, we constructed libraries of pegRNAs designed to generate all possible nucleotide changes, as well as generating one additional synonymous substitution in the *EGFR* gene’s coding DNA sequence (CDS), in the region spanning exon 18 to exon 24 (Syn-exon18 through Syn-exon24, Supplementary Tables 1 and 2). Since PC-9 cells harbor a 15-bp in-frame deletion in exon 19 of *EGFR*, pegRNAs targeting this region were designed based on the deleted sequence. To achieve a high frequency of prime editing, we initially calculated the DeepPrime-FT score (DeepPrime-FT is a fine-tuned model for predicting the activity of PE4max used with an optimized scaffold in HEK293T cells)^31^ for pegRNAs designed to generate all possible 1-bp substitutions in the targeted region. Subsequently, we excluded all pegRNAs with left homology arms (LHAs) that were shorter than 5 bp and chose the three pegRNAs with the highest DeepPrime-FT scores for each SNV. We also introduced additional 1-bp synonymous substitutions in codons different from those harboring the intended substitutions in the LHA of the pegRNA RT template. This manipulation was done to maintain the minimal length of the right homology arm (RHA) of the RT template, which is crucial for efficient prime editing^31^. These additional substitutions serve multiple purposes: (1) Sequencing reads can be filtered based on these additional substitutions to distinguish true edited reads from reads containing sequencing errors or partial incorporation of the RT template. (2) The incorporation of additional mismatches near the intended substitution increases the prime editing efficiency by weakening the repair of the resulting heteroduplex^34^ and thereby could bypass MMR. Out of 2,610 possible SNVs, 95 were excluded from the library because pegRNAs could only be designed at distances exceeding the maximum RT template length specified by the DeepPrime model, resulting in a total 2,515 SNVs. In pegRNAs targeting 36 of a total of 2,515 SNVs (1.4%), additional synonymous substitutions were not introduced due to unavailable silent substitution sites. As negative controls, we included 198 pegRNAs with spacer sequences identical to those used in SNV-inducing pegRNAs, along with RT templates that matched the reference sequence and were designated as “sham-editing” pegRNAs. Additionally, we incorporated 198 pegRNAs that do not target any sequence in the human genome; these were labeled as “nontargeting” pegRNAs and had RT templates of randomly annotated context sequences. In total, we designed a total of 7,884 pegRNAs (Supplementary Tables 1 and 2).

For the Syn-RPL15 library, we calculated the DeepPrime-FT scores for pegRNAs designed to generate all possible 1-bp substitutions within the CDS of exon 2 of *RPL15*. We excluded pegRNAs with short LHAs and chose the three pegRNAs with the highest DeepPrime-FT scores, as we did when designing the syn-Exon18 through syn-Exon24. Subsequently, an additional 1-bp synonymous substitution was introduced in the LHA of the RT template. We designed a total of 1492 pegRNAs targeting 500 SNVs in *RPL15* (Supplementary Tables 1 and 2).

To design a library of pegRNAs designed to generate all possible nucleotide changes in the CDS of exon 20 of *EGFR* for NRCH-PEmax (NRCH-exon20), we calculated the DeepPrime-FT score from a pilot version of the original model (specific for NRCH-PE2max in HEK293T cells) for pegRNAs designed to generate all possible 1-bp substitutions in the targeted region. We chose the three pegRNAs with the highest DeepPrime-FT score for each SNV. As negative controls, 86 sham-editing pegRNAs and 85 nontargeting pegRNAs were included. In total, we designed a total of 1,845 pegRNAs (Supplementary Tables 1 and 2).

#### Construction of pegRNA libraries and cloning

The pegRNA library targeting *EGFR* was divided into subsets of exon-specific libraries (Syn-exon18 through Syn-exon24, Supplementary table 2). Each subset included sham-editing pegRNAs with spacer sequences used in the subset and the same number of nontargeting pegRNAs.

Pooled 228-bp oligonucleotides for plasmid construction were array-synthesized by Twist Bioscience. For Syn-exon18 through Syn-exon24 and Syn-RPL15, oligonucleotides were designed to include the following elements (Extended Data Fig. 1b): (1) a 19-bp homology arm with a 3’ terminus from a U6 promoter; (2) a 20-bp sequence with a G at the 5’ terminus, followed by a 19-bp spacer sequence; (3) a random 18-bp sequence flanked by a BsmBI cut site on either side (11 bp each); (4) a RT template and a primer binding site (PBS); (5) an 8-bp linker sequence obtained from pegLIT tools ^62^ followed by a 37-bp tevopreQ1 and a 7-bp poly-T sequence; (6) an 18-bp unique barcode sequence corresponding to a unique pegRNA sequence, which contained a constant “GTCAG” sequence for subsequent analysis; (7) a random “Buffer” sequence of various lengths added to pegRNAs with shorter RT templates and PBSs to ensure that all oligonucleotides were of equal length; and (8) a 20-bp unique homology arm for subset-library amplification.

#### Unique homology arms for *EGFR*; 5’-3’

*EGFR* exon 18: TGCGACCGTAATCAAACCAA *EGFR* exon 19: ATGCAATCGGCCTGGTATTT *EGFR* exon 20: AAATGGATGCCTTGTGCGAA *EGFR* exon 21: AAACGGAGCCATGAGTTTGT *EGFR* exon 22: AAACTGGAGGCGGCAAATTA *EGFR* exon 23: ACCGCGCTCGAAGAATTTAA *EGFR* exon 24: TCCTCAGCCGATGAAATTCC

Oligonucleotides (∼4 ng) were amplified using primer set 1/2 and Q5 High-Fidelity DNA polymerase (NEB). Unique reverse primers contained an 8-bp random Unique Molecular Identifier (UMI) sequence^24^ for pegRNA-based analysis (Supplementary Table 8). PCR cycling conditions were as follows: 30 s at 98℃, 30 s at 68℃, 2 min at 72℃, for 14 cycles. The resulting amplicons were size-selected by electrophoresis on a 2% agarose gel and assembled into BsmBI-linearized Lenti-gRNA-Puro^63^ (Addgene, 84752) using NEBuilder HiFi DNA Assembly Master Mix (NEB). After the assembly reaction, the product was isopropanol precipitated and electroporated into Endura electrocompetent cells (Lucigen) to generate a scaffoldless plasmid library.

An Improved version of the SpCas9 sgRNA scaffold sequence^64^ (Supplementary Table 8) and BsmBI cut sites were synthesized, PCR-amplified, and ligated into the TOPO vector (Invitrogen, Zero Blunt TOPO PCR cloning kit). A ligation reaction was performed using the BsmBI-digested scaffoldless plamid library generated above and the BsmBI-digested sitcky-ended sgRNA scaffold sequences. The reaction mixtures were pooled, isopropanol precipitated, and electroporated into Endura electrocompetent cells to construct the final plasmid library.

#### Cell lines and culture

PC-9 (RRID: CVCL_B260) cells were purchased from Riken Cell Bank. PC-9 cells were cultured in RPMI 1640 (Gibco, 2.05mM L-glutamine) supplemented with 10 mM HEPES, 1 mM sodium pyruvate with 10% FBS (RDT) and 1% penicillin-streptomycin (Gibco). Cells were passaged every 3 days and were maintained in 37℃ incubators with 5% CO_2_. HEK293T cells (ATCC) were cultured in DMEM (Gibco) with 10% FBS (RDT) at 37℃ with 5% CO_2._ The doses of polybrene, puromycin, and blasticidin were as follows: 8 μg/ml, 0.2 μg/ml, and 0.3 μg/ml, respectively.

#### Construction of plasmid vectors

All primers used for cloning are listed in Supplementary Table 8. To generate a lentiviral vector for expressing PEmax, we assembled three linear DNA fragments: (1) a linearized lentiviral backbone from Lenti-Cas9 Blast (Addgene #52962)^65^ (XbaI and EcoRI (NEB); (2) a sequence encoding Cas9 containing R221K, N394K and H840A mutations amplified from pCMV-PEmax-p2A-BSD^34^ (Addgene #174821) using primer pair 4/5; and (3) optimized MMLV-RT and NLS sequences and a blasticidin resistance gene amplified from pCMV-PEmax-p2A-BSD using primer pair 6/7. The assembled vector is referred to as pLenti-PEmax-p2A-BSD.

To generate a lentivirus for expressing NRCH-PEmax, we assembled five linear DNA fragments: (1) a linearized lentiviral backbone from Lenti-Cas9 Blast (Addgene #52962) digested with XbaI and EcoRI (NEB); (2) a sequence encoding the N-terminal region of Cas9 containing R221K and N394K mutations amplified from pCMV-PEmax-p2A-BSD^34^ (Addgene #174821) using primer pair 8/9; (3 and 4) sequences encoding the C-terminal region of Cas9-NRCH amplified with primers containing H840 sequences (primer pair 10/11 and 12/13) from pCMV-Cas9-NRCH^32^ (Addgene #136926); and (5) optimized MMLV-RT and NLS sequences amplified from pCMV-PEmax-p2A-BSD^34^ (Addgene #174821) using primer pair 14/15. The assembled vector is referred to as pLenti-Cas9-NRCH-PEmax-p2A-BSD.

#### Lentivirus production

HEK293T cells were seeded in 150-mm culture dishes at a density of 1×10^7^ cells per dish 18 hours before transfection. Transfection was performed using lipofectamine 3000 (Invitrogen) transfection reagent according to the manufacturer’s protocol. In brief, transfer plasmids containing the gene of interest, psPAX2, and pMD2.G were mixed at a molar ratio of 4.3:3.4:1.9 pmol and diluted into 1 ml of Opti-MEM (Life Technology). P3000 reagent was added to the diluted plasmid mixtures such that the ratio of μg DNA: μL P3000 reagent was 1:2. Lipofectamine 3000 reagent was diluted into 1 ml of Opti-MEM such that the ratio of μg DNA: μL lipfectamine 3000 reagent was 1:2.25, and the resulting solution was added to the DNA mixture. The mixture was incubated for 20 min and added to cells. At 6 h after transfection, 20 ml of growth medium was added to refresh the cells.

The growth medium was harvested 24 and 52 h after transfection. The harvested medium was centrifuged at 2,000g for 10 min to pellet cell debris. The supernatant was filtered through a Millex-HV 0.45-μm low protein-binding membrane (Millipore), divided into aliquots, and kept frozen at -80℃ until use.

#### Generation of cell lines

To generate prime editor-expressing cells, wild-type PC-9 cells were transfected with a mixture of a plasmid expressing SpCas9 (pRGEN-Cas9-CMV/T7-Puro-RFP; purchased from ToolGen, Korea) and two plasmids expressing sgRNAs (pRG2; Addgene #104174)^67^. The sgRNA sequences were as follows:

MLH1-sgRNA1: 5’-TGGAGGGGGATACAACAAAG(GGG)-3’ MLH1-sgRNA2: 5’-GCCTGGGGCTGACTGGACAG(GGG)-3’

PC-9 cells were electroporated using an NEPA21 electroporator (NepaGene) following the manufacturer’s protocol. In brief, 5 μg of SpCas9-expressing plasmid and 2.5 μg of each of two sgRNA-expressing plasmids were mixed and electroporated with a 7.5 ms pulse length at 125V of poring voltage. Cells expressing the transgene were sorted using FACS at day 5 post-transfection (Supplementary Fig. 6) and subjected to deep sequencing using primer pair 22/23 (Supplementary Table 8).

The resulting PC-9-MLH1KO cell line was subsequently transduced with lentiviral vectors expressing PEmax or NRCH-PEmax and the blasticidin resistance gene, as described above. The day after transduction, cells were refreshed with growth medium containing 3 μg ml^-1^ of blasticidin (Invivogen). After 2 weeks of selection, the obtained PEmax-expressing MLH1-knockout PC-9 cells were aliquoted and used for further experiments.

To generate T790M-expressing PC-9 cells, PEmax-expressing MLH1-knockout PC-9 cells were transfected with a mixture of a plasmid expressing BE4max (pCMV-BE4max; Addgene #112093)^68^ and a plasmid expressing sgRNA (pLKO5.sgRNA.EFS.tRFP; Addgene #57823)^69^. The sgRNA sequence was as follows:

T790M-sgRNA : 5’-ATCACGCAGCTCATGCCCTT(CGG)-3’

PEmax-expressing MLH1-knockout PC-9 cells were electroporated as previously described with 7.5 μg of BE4max-expressing plasmid and 2.5 μg of sgRNA-expressing plasmid. Cells expressing the transgene were sorted using FACS at day 7 post-transfection.

#### SynPrime pooled assay

For all experiments, cells were transduced in two biological replicates for each exon. Twenty-four hours before transduction of the lentiviral library, cells were seeded to achieve a representation of approximately 10,000 cells per pegRNA. Transductions were performed at a multiplicity of infection (MOI) of 0.5, ensuring that every pegRNA was represented in approximately 5,000 cells. After 24 h of infection, the medium was replaced with medium containing puromycin, and the cells were cultured for an additional 9 days. Upon puromycin removal (day 0), cells were harvested in quantities sufficient for 5,000-fold coverage of the pegRNA libraries. The remaining cells were split into a drug arm and an untreated arm and were maintained in quantities sufficient for 10,000-fold coverage of the pegRNA libraries. Drugs were used at the following concentrations: afatinib (Santa-Cruz Biotechnology, sc-364398) at 3 nM and Osimertinib (SelleckChem, S7297) at 8 nM. The cells were passaged every 3 days for an additional 10 days and collected for genomic DNA extraction.

For RPL15 evaluations, cells were transduced as previously described. After 24 hours of infection, the medium was replaced with medium containing puromycin, and the cells were cultured for an additional 4 days. Upon puromycin removal (day 0), cells were harvested in quantities sufficient for 5,000-fold coverage of the pegRNA libraries. The remaining cells were maintained in quantities sufficient for 10,000-fold coverage of the pegRNA libraries. The cells were passaged every 3 days for an additional 10 days and collected for genomic DNA extraction.

#### Genomic DNA preparation and deep sequencing

Genomic DNA was extracted using a Wizard Genomic DNA Purification Kit (Promega) according to the manufacturer’s protocol. Using the isolated genomic DNA as template, endogenous target genomic regions or RT template, pegRNA-specific barcodes and UMI (Extended Data Fig. 1b) were amplified and prepared for deep sequencing through two rounds of PCR using 2X Pfu PCR Smart Mix (Solgent). In the first step, genomic DNA was divided into multiple 50-μL reactions containing 3 μg of genomic DNA, 20 pmol of forward primer, 20 pmol of reverse primer (primers 24 to 39 for endogenous target genomic regions and primers 40 to 45 for RT template, pegRNA-specific barcodes and UMI, Supplementary Table 8), and 25 μL of PCR pre-mix. The PCR cycling parameters were as follows: an initial 2 minutes at 95 °C, followed by 30 s at 95 °C, 30 s at 60 °C, and 40 s at 72 °C, for 20 cycles, and a final 5-min extension at 72 °C. The total amount of genomic DNA for each experiment represented more than 10,000× coverage of the library, assuming 6.6 μg of genomic DNA per 10^6^ cells ^70^. Amplicons for each experiment were pooled and concentrated with a MEGAquick-Spin Total Fragment DNA Purification kit (iNtRON Biotechnology) and size-selected with agarose gel electrophoresis.

In the second PCR step, a total of 20 ng of purified PCR product from the first step was used in two separate 50-μL reactions with 20 pmol of Illumina indexing primers in each reaction (primer pair 46/47). The PCR cycling parameters were same as in the 1^st^ PCR except for the number of PCR cycles; the 2^nd^ PCR was performed for 5 cycles. Amplicons for each experiment were size-selected with agarose gel electrophoresis and sequenced using a NovaSeq 6000 (Illumina).

#### Data analysis and variant filtering

To identify SNVs introduced into the cells transduced with the Syn-RPL15 and Syn-exon18 through Syn-exon24 libraries, a SNV reference sequence library was created. The library was generated from the CDS and 5 nucleotides of adjacent intronic sequence (NM_005228.5). The sequences included in this reference library only contained the intended SNV and the additional synonymous substitution, without any other mismatches or indels. For the NRCH-exon20 library, a SNV reference sequence library was created, which contained only the intended SNVs without any other mismatches or indels. Subsequently, the processed reads from targeted deep sequencing were aligned to this SNV reference sequence library, and counts were recorded only when the next-generation sequencing (NGS) reads exhibited a perfect match to the SNV sequence library. Unedited wild-type reads were also identified when the NGS reads perfectly matched the reference sequence (NM_005228.5).

To distinguish true prime-edited reads from errors generated by partial RT template incorporation or errors occurring during NGS or library preparation, we calculated the odds ratio (OR) and *P*-value using Fisher’s exact test between NGS reads from cells at day 0 and NGS reads from unedited cells as follows:

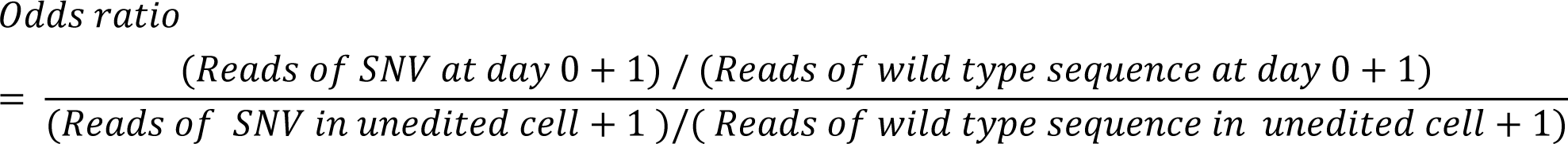

After calculating the OR and *P*-value of each SNV, we identified true edited SNVs as those with a *P*-value < 0.05 and an OR > 3. Additionally, we further removed SNVs that had fewer than 0.5 read per million (RPM) in the cells at day 0. These criteria help ensure that the identified SNVs are statistically significant and have a sufficient level of representation in the sequencing data.

#### Calculation of adjusted LFCs

For syn-RPL15 data, we calculated the log_2_ fold change (LFC) of each SNV by comparing allele frequencies in the cells at day 10 with those in the cells at day 0. For Syn-exon20 data, we calculated the LFC of each SNV by comparing allele frequencies in the untreated cells at day 10 with those in the cells at day 0 for EGFR dependency analysis (Fig. 3g-i). Additionally, we calculated the LFC of each SNV by comparing the allele frequencies in the drug-treated cells at day 10 with those in the untreated cells at day 10 samples to generate a drug resistance profile. These LFC values for each SNV were normalized using LOWESS (Locally Weighted Scatterplot Smoothing) regression.

#### Modelling positional biases of data integration and normalizing across exons

To account for the effect of the sequence context on the editing efficiency, we normalized the LFC of each SNV using the LFC of synonymous SNVs, which are considered to have a neutral LFC as they do not introduce any amino acid change. We calculated the regressed synonymous SNV LFC at each position in each exon using LOWESS regression.

Subsequently, we obtained standardized LFC values by subtracting the synonymous SNV LFC at each position (LOWESS regressed) and dividing by the standard deviation of synonymous SNVs in each exon for comparison between exons (Extended Data Fig. 2b). Next, we averaged the standardized LFC of each replicate to obtain the adjusted LFC^21^.

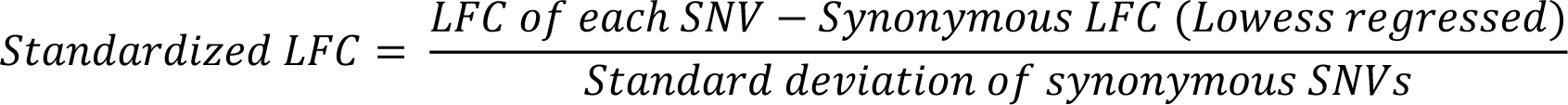

To calculate the adjusted LFC based on protein variants, we averaged the adjusted LFC values of SNVs inducing the same protein variant.

#### SNV drug resistance classifications

We analyzed drug resistance induced by the Syn-exon18 through Syn-exon24 libraries using cutoff values determined by the distribution of resistance scores of synonymous SNVs within each exon. Functional classifications were made as follows:

- Resistance: SNV resistance scores that were above the 99.7th percentile relative to scores of the synonymous SNVs in both replicates.
- Sensitive: SNV resistance scores that were under the 95th percentile relative to scores of the synonymous SNV in both replicates.
- Intermediate: SNVs fell into neither the ‘Resistance’ nor ‘Sensitive’ categories.

To obtain a classification based on protein variants, we calculated the LFC of protein variants that could be induced by multiple SNVs, as previously described. The classification criteria used in each dataset were the same as those employed in the SNV-based classification.

#### pegRNA barcode count and library-based screening analysis

The overall method of counting pegRNA barcodes and calculating the fold change of SNV frequencies between treatment and control groups, such as cells at day 10 from the afatinib treated arm and cells at day 0, was performed as described previously using an in-house Python code, which was also used in our previous study (see ‘Code Availability’)^24^. In our deep sequencing analysis of pegRNAs, we identified a barcode motif, permitting a single base mismatch, and a 20-bp unique homology sequence, also allowing for one base mismatch. Subsequently, we defined the UMI sequence as the motif found between the barcode and the 20-bp unique homology sequence. UMIs were then collapsed if the fold change in read count was three or greater between two UMIs with a single base mismatch. These collapsed UMIs were categorized into four distinct groups for each pegRNA. Given that a single SNV can be induced by two or three pegRNAs, we conducted the MAGeCK test using eight to twelve pegRNA-UMI group pairs. In this analysis, pegRNAs were treated as sgRNAs, and SNVs were considered to be gene names. MAGeCK version 0.5.9.3^43^ was used to calculate the fold change of SNV frequencies between treatment and control groups. Then, we normalized the LFC of each SNV using the LFCs of nontargeting and sham-editing pegRNAs. In the pegRNA-based screening analysis, we also filtered out SNVs according to the criteria described in “Data analysis and variant filtering”. The remaining pegRNAs were then classified depending on their normalized LFCs in both replicates using cutoff values determined by the distribution of the LFCs of the non-targeting and sham-editing pegRNAs in each exon. Classification thresholds were the same as we described in “SNV drug resistance classifications”.

#### Resistance profiles of EGFR variants in previously published literature

We identified 16 SNVs within the *EGFR* gene from the ClinVar database^71^, all of which were classified as having clinical significance related to drug response. Additionally, for SNVs classified as conferring resistance to afatinib, osimertinib in the absence of T790M, or osimertinib in the presence of T790M in at least one dataset, we manually verified whether they had been previously reported in the PubMed database (Supplementary Fig. 4c).

#### Evaluation of resistance in conventional manner

PegRNA sequences designed to generate K754Q, G930R, and E931K SNVs were individually cloned into BsmBI-linearized Lenti gRNA Puro (Addgene #87245). The following 4 fragments were assembled with the linearlized vector by Golden Gate cloning: (1) annealed spacer oligonucleotides with overhangs for cloning; (2) an improved SpCas9 sgRNA scaffold sequence with overhangs for cloning (primer 3); (3) an annealed epegRNA RT template/PBS 3’ extension with overhangs for cloning; and (4) tevopreQ1 and poly T sequences with overhangs for cloning.

In total, 3 million PEmax-expressing MLH1-knockout PC-9 cells per pegRNA were seeded in 150-mm culture dishes 24 hours before transduction in triplicate. The cells were infected with lentivirus harboring pegRNA sequences at a high MOI (∼1). For a negative control, a separate population of 3 million cells were infected with lentivirus harboring empty vectors. The day after transduction, the medium was replaced with fresh medium containing puromycin, after which cells were incubated for an additional 9 days.

Ten days after infection, cells transduced with pegRNA sequences and cells transduced with empty vector were mixed at a 25:75 ratio. These mixed cell populations were then divided into untreated and drug-treated conditions. Drugs were used at the following concentrations: afatinib at 3 nM and osimertinib at 12 nM. The mixed cell populations were cultured for 10 days and subsequently subjected to deep sequencing. Each pegRNA-targeted genomic site was amplified using site-specific primers.

#### Statistical analysis

All statistical tests described were performed as two-tailed tests using Python software packages.

#### Data visualization

Figures were created with Python. Schematics were created with BioRender.com. PyMOL (version 2.5.5) was used to map the variants onto the following crystal structures from the Protein Data Bank: PDB: 6JXT, (EGFR complex with osimertinib).

#### Data availability

We have submitted the deep sequencing data from this study to the National Center of Biotechnology Information’s Sequence Read Archive under accession number PRJNA1018283. We have provided the datasets used in this study as Supplementary Tables 2–5.

#### Code availability

Screening data were analyzed with in-house custom Python scripts and MAGeCK (version 0.5.9.3). Custom Python scripts (version 3.8.16) were used to generate input files based on grouped UMIs for MAGeCK. They are available at https://github.com/oreolic/SGE_EGFR.

---

**Extended Data Figure 1.**
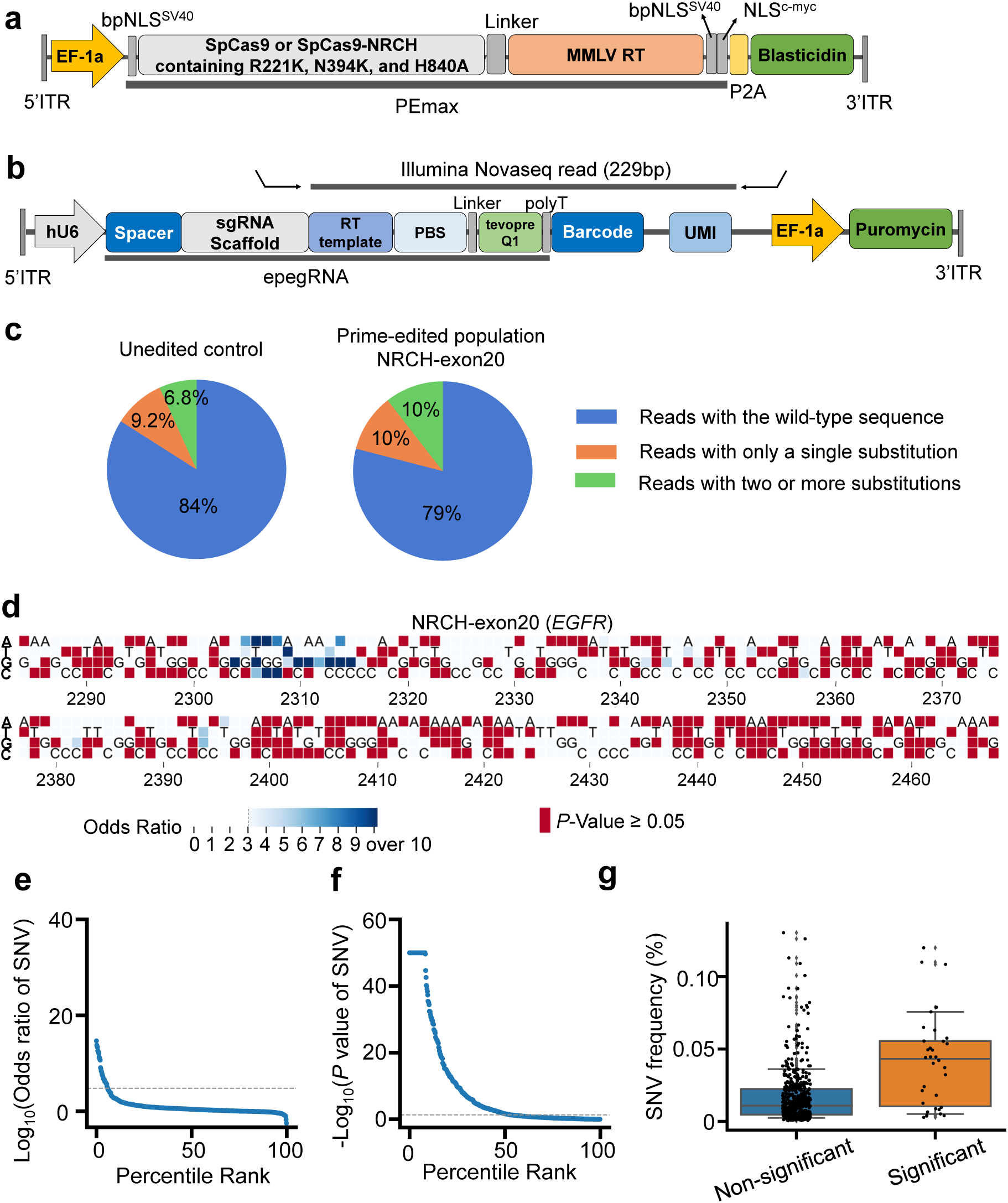
Sequencing errors can hinder the accurate identification of SNVs induced by prime editing. **a,b**. Maps of lentiviral vectors used for the expression of prime editor (PEmax) (a) and pegRNAs (b). bpNLS, bipartite nuclear localization signal; MMLV-RT, human codon-optimized Moloney murine leukemia virus reverse transcriptase; ITR, inverted termainal repeat. (b) Locations of PCR primers used to deep-sequence a 229-bp region containing the RT-template, PBS (primer binding site), pegRNA barcode, and unique molecular identifier (UMI) are shown. **c**, Proportion of sequencing reads containing substitution(s) in unedited PC-9 cells expressing NRCH-PEmax (left, unedited control) and in those ten days after transduction with the NRCH-exon20 library (right) for editing exon 20 of *EGFR*. **d**, Heatmap showing odds ratios and/or *P*-values of 558 (= 186 x 3) SNVs generated by prime editing in exon 20 of *EGFR* ten days after the transuction of the NRCH-exon20 library. SNVs with *P*-values greater than 0.05 by the two-sided Fisher’s exact test are indicated in red; in these cases, odds ratios are not shown. SNVs with odds ratios lower than 3 are shown in white background. The numbers at the bottom of each heatmap represent the location in the *EGFR* coding sequence. At each position, the nucleotide in the reference sequence is shown. **e**,**f**, Distribution of odds ratios (e) and *P*-values (f) of 558 SNVs in cells transduced with the NRCH-exon20 library. The dashed horizontal lines indicate the position at which the odds ratio = 3 (e) and the *P*-value = 0.05 (f). **g**, Distribution of observed SNV frequencies in PC-9 cells expresing PEmax ten days after transduction with NRCH-exon20. The number of SNVs *n* = 524 (Nonsignificant), *n* = 34 (Significant). Boxes represent the 25th, 50^th^, and 75th percentiles, and whiskers show the 10th and 90th percentiles.

**Extended Data Figure 2.**
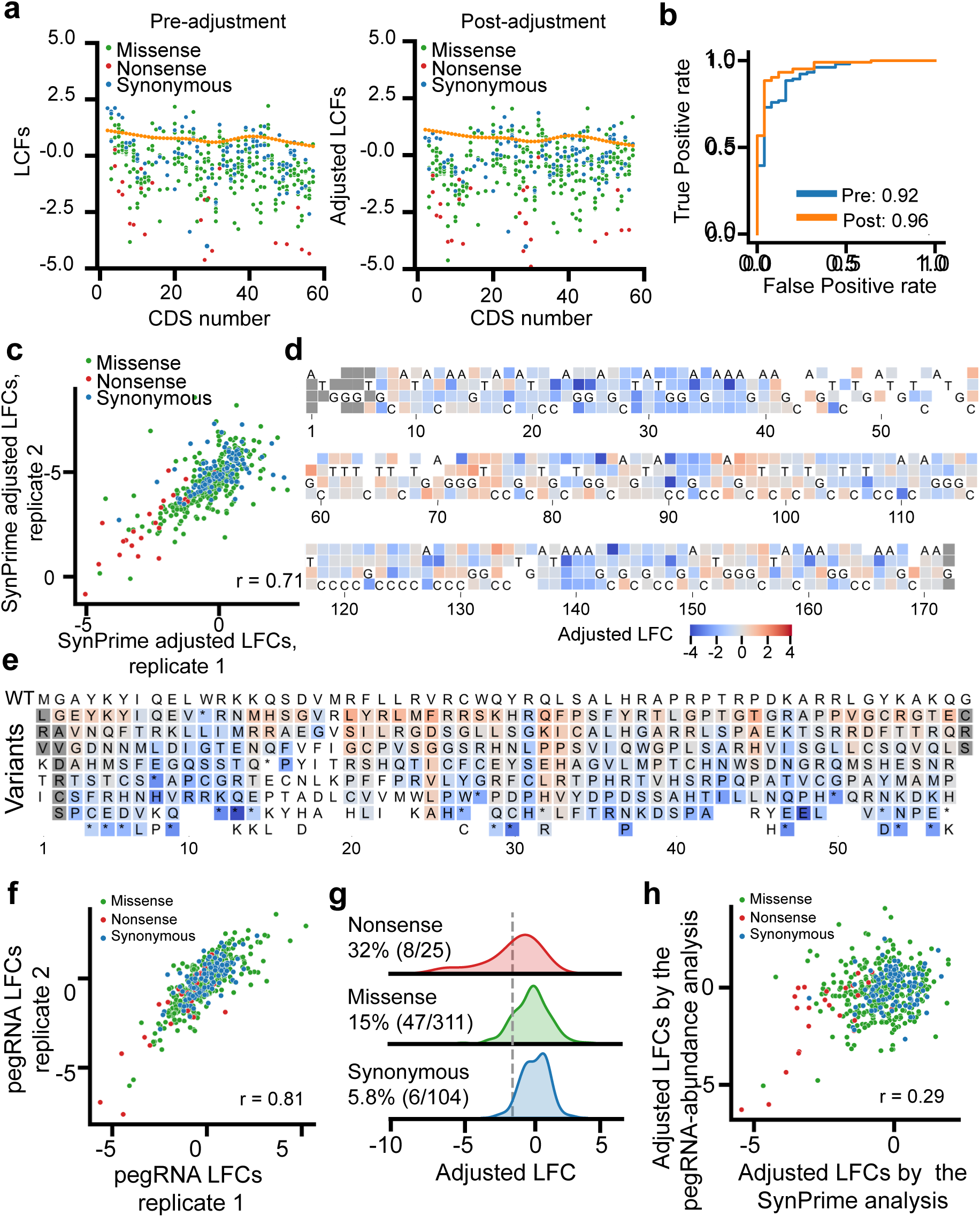
Optimization of SynPrime and comparison with pegRNA-based analysis. **a**, Correction of positional biases. To correct positional biases in LFCs, we employed LOWESS regression by utilizing synonymous SNVs, which were assumed to have no functional effect. The LOWESS regression curves are shown as orange lines. The overall depletion of nonsense SNVs was more distinct after the adjustment. **b**, ROC-AUC analysis to determine the effect of positional bias correction for sets of nonsense (the number of SNVs *n* = 25) versus synonymous SNVs (*n* =104) in exon 2 of *RPL15*. Pre, ROC-AUC before adjustment; Post, ROC-AUC after adjustment. **c**, Correlation between SynPrime LFC values from two biological replicates. Pearson correlation coefficients (r) are shown. The number of SNVs *n* = 440. **d,e**, Heatmaps showing adjusted LFCs of 516 (= 172 x 3) SNVs (d) and 336 protein variants (e) generated by prime editing in exon 2 of *RPL15*. SNVs (d) and protein variants (e) with *P*-values greater than 0.05 or odds ratios lower than 3 were excluded from the analysis and are shown in white background. SNVs (d) and protein variants (e) for which no pegRNAs were designed are shown as gray boxes. The numbers at the bottom of each heatmap represent the location in the *RPL15* coding sequence (d) and in the RPL15 amino acid sequence (e). At each position, the nucleotide (d) and amino acid (e; WT, wild-type; top) in the reference sequences are shown. **f**, Correlation between adjusted LFC values of SNVs determined by pegRNA abundance-based analysis from two biological replicates. Pearson correlation coefficients (r) are shown. The number of SNVs *n* = 440. **g**, Kernel density estimation plots of adjusted LFCs of SNVs in *RPL15* determined by pegRNA abudance-based analysis as a function of the SNV category. For each category, the number and percentage of SNVs with adjusted LFC values lower than a cutoff value (the gray dashed vertical line), representing the 5th percentile of the adjusted LFC values of synonynous mutations, are shown. **h**, Correlation between adjusted LFC values of SNVs calculated from SynPrime evaluations and those from pegRNA abundance-based analysis. Pearson correlation coefficients (r) are shown. The number of SNVs *n* = 440.

**Extended Data Fig. 3.**
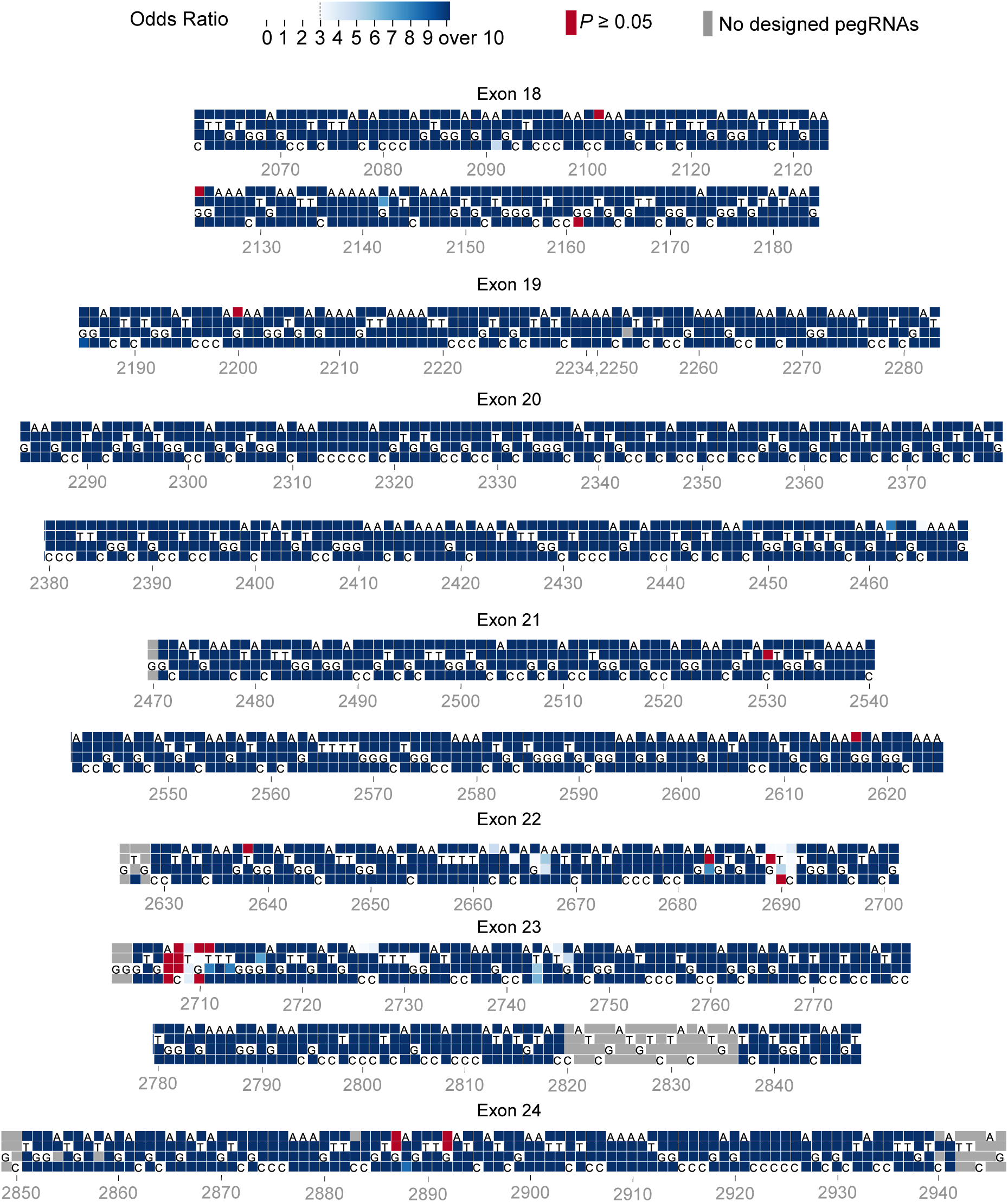
Identified SNVs. Heatmap showing odds ratios and/or *P*-values of 2,610 (= 870 x 3) SNVs generated by prime editing in exons 18-24 of *EGFR* ten days after the transduction of the Syn-exon18, Syn-exon19, …, Syn-exon24 libraries. SNVs with *P*-values greater than 0.05 by the two-sided Fisher’s exact test are indicated in red; in these cases, odds ratios are not shown. SNVs with odds ratios lower than 3 are shown in white background. The numbers at the bottom of each heatmap represent the location in the *EGFR* coding sequence. At each position, the nucleotide in the reference sequence is shown. Edited reads were identified based on the the presence of both the intended edit and an additional synonymous edit.

**Extended Data Fig. 4.**
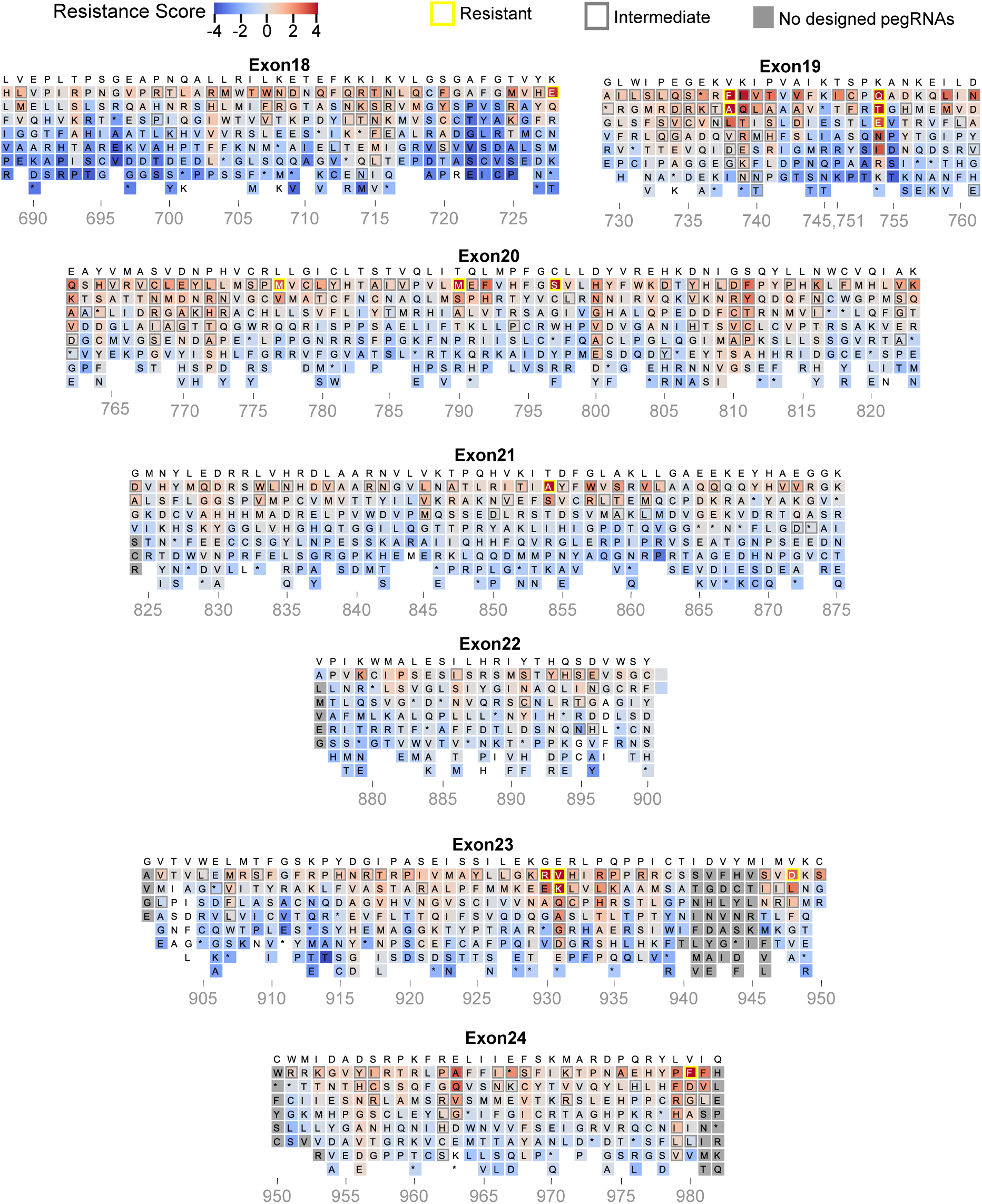
Heatmap showing afatinib resistance scores of 1,817 protein variants. These variants were generated by prime editing in exons 18-24 of *EGFR* in PC-9 cells. Boxes outlined in yellow and gray indicate protein variants causing resistant and intermediate phenotypes, respectively. The numbers at the bottom of each heatmap represent the location in the EGFR amino acid sequence. At each position, the amino acid in the reference sequence is shown at the top. Thirty protein variants with *P*-values greater than 0.05 or odds ratios lower than 3 were excluded from the analysis and are shown in white background. Protein variants for which no pegRNAs were designed are shown as gray boxes.

**Extended data figure 5.**
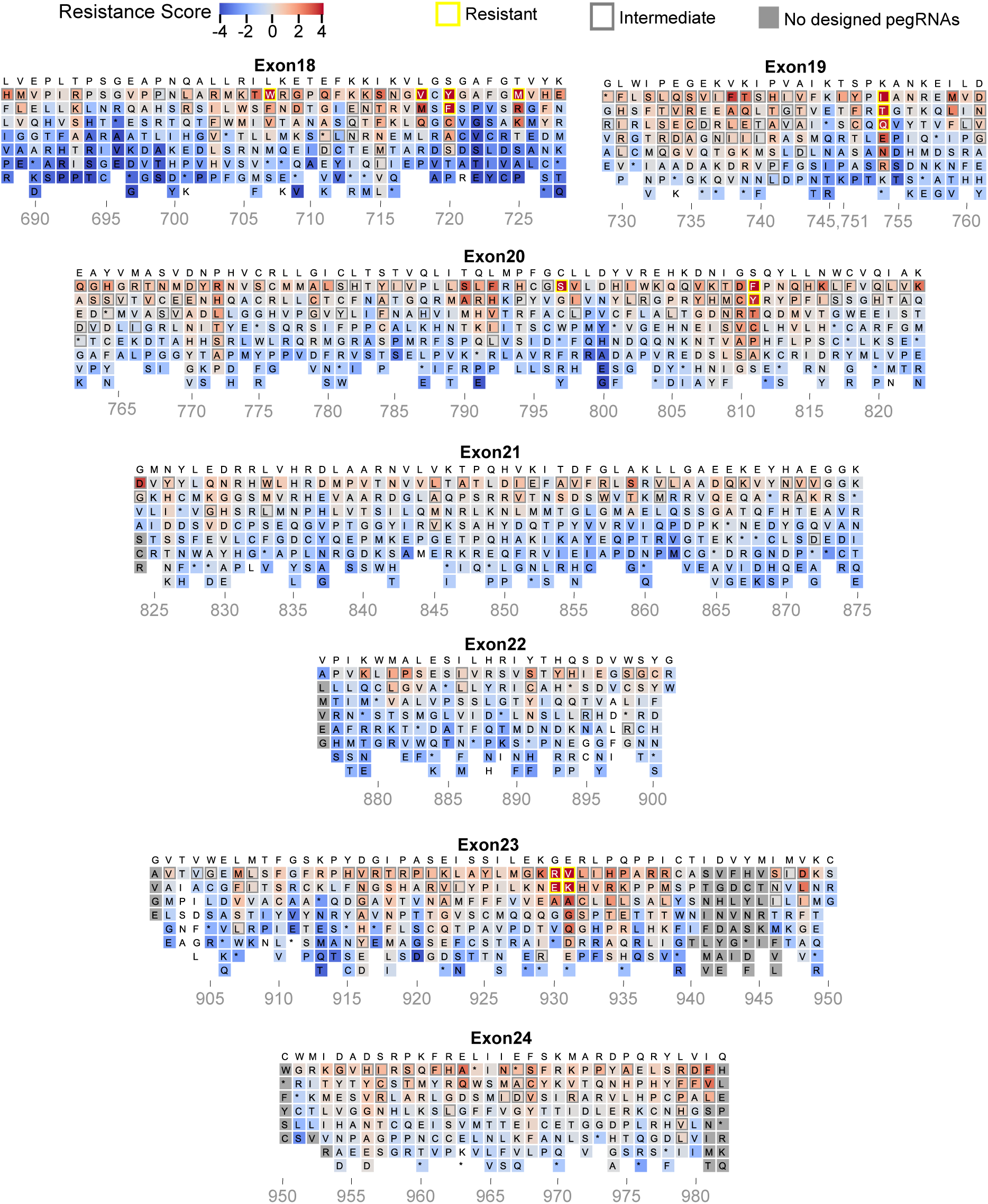
Heatmap showing osimertinib resistance scores of 1,817 protein variants. These variants were generated by prime editing in exons 18-24 of *EGFR* in PC-9 cells. Boxes outlined in yellow and gray indicate protein variants causing resistant and intermediate phenotypes, respectively. The numbers at the bottom of each heatmap represent the location in the EGFR amino acid sequence. At each position, the amino acid in the reference sequence is shown at the top. Thirty protein variants with *P*-values greater than 0.05 or odds ratios lower than 3 were excluded from the analysis and are shown in white background. Protein variants for which no pegRNAs were designed are shown as gray boxes.

**Extended Data Fig. 6.**
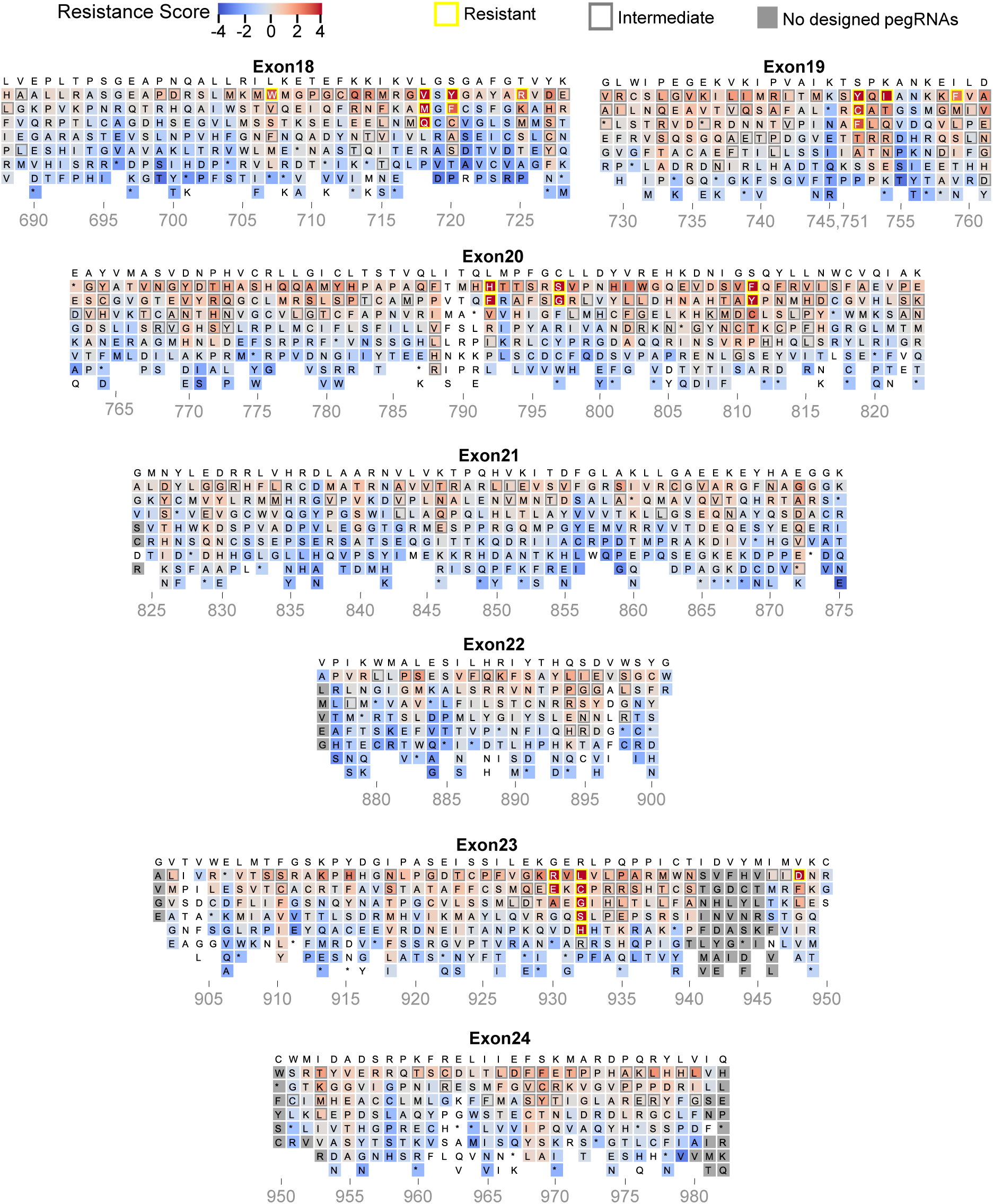
Heatmap showing osimertinib resistance scores of 1,817 protein variants in the presence of a co-occuring T790M mutation. These variants were generated by prime editing in exons 18-24 of *EGFR* in PC-9 cells containing the T790M mutation. Boxes outlined in yellow and gray indicate protein variants causing resistant and intermediate phenotypes, respectively. The numbers at the bottom of each heatmap represent the location in the EGFR amino acid sequence. At each position, the amino acid in the reference sequence is shown at the top. Ninety-four protein variants with *P*-values greater than 0.05 or odds ratios lower than 3 were excluded from the analysis and are shown in white background. Protein variants for which no pegRNAs were designed are shown as gray boxes.

